# Coffee consumption affects adipose tissue remodeling in a high-fat diet-induced obesity model by different mechanisms in rats

**DOI:** 10.1101/2025.07.18.665488

**Authors:** Marilia Hermes Cavalcanti, Teresa Helena Macedo da Costa, Amílcar Sabino Damazo, Sandra Fernandes Arruda

## Abstract

This study investigated the effects of coffee consumption on thermogenesis and lipid metabolism in brown (BAT), inguinal (iWAT), and epididymal (eWAT) adipose tissues, and their relationship with inflammatory and redox responses in rats. Wistar rats were fed control (CT), high-fat (HF), control with coffee (CT+), or high-fat with coffee (HF+) diets. The HF diet increased the adipocyte area in all adipose tissues, hepatic cholesterol and triglyceride levels, and liver steatosis scores. In BAT, it increased the number of unilocular adipocytes; upregulated *Ppargc1a*, *Ucp1*, *Vegfa*, *Vegfr2,* and *Il10* mRNA levels; and downregulated *Acaca* and *Fasn* mRNA levels. In iWAT, it elevated *Vegfr2* and *Slc2a4* mRNA levels and carbonyl content, and reduced lipid peroxidation and *Ucp1* mRNA and protein levels. In eWAT, it upregulated *Slc2a4,* and downregulated *Acaca*, *Fasn, Prkaa1,* and *Prkaa2* mRNA levels. The HF+ diet reduced the adipocyte area; unilocular adipocytes; and *Acaca, Fasn,* and *Il10* mRNA levels in BAT. Coffee decreased catalase activity in BAT independent of diet type. In iWAT, the HF+ diet reduced adipocyte area and glutathione peroxidase (GPx) activity, while increasing the number of multilocular adipocytes, *Prdm16* and *Vegfr2* mRNA levels, and UCP1 protein. Catalase activity decreased with coffee intake, regardless of diet type. In eWAT, coffee increased multilocular adipocytes and *Prdm16* mRNA levels, and decreased *Fasn* mRNA, independent of diet type. Coffee combined with the control diet increased *Ppargc1a* and liver steatosis scores, while decreasing *Acaca* and *Vegfr2* mRNA levels and catalase and GPx activities compared to those with the control diet. Overall, coffee consumption at levels comparable to the typical Brazilian intake partially mitigated BAT whitening, likely through the modulation of lipogenic and lipolytic gene expression. In the iWAT, coffee had a more pronounced effect by enhancing thermogenic and angiogenic gene expression, suggesting potential molecular mechanisms through which coffee may aid in obesity management.

## Introduction

Recently, the prevalence of metabolic disorders, particularly obesity, has increased at an alarming rate and became a major public health challenge worldwide [1]. Excess adipose tissue is a well-established risk factor for type 2 diabetes, hypertension, and cardiovascular disease [2]. This condition is characterized by low-grade chronic inflammation, which contributes to oxidative stress and ultimately leads to cellular and molecular damage that exacerbates metabolic dysfunction. Therefore, “browning” of white adipose tissue is a promising therapeutic approach. This process involves the phenotypic conversion of energy-storing white adipocytes into beige adipocytes with a thermogenic capacity similar to that of brown adipocytes, thereby increasing the overall energy expenditure and reducing lipid accumulation [3,4].

Certain dietary components can stimulate brown adipose tissue (BAT) activity and promote the transdifferentiation of white adipocytes into beige cells in high-fat diet-induced obesity models. These effects not only contribute to weight loss but also trigger adaptive remodeling of adipose tissue, including the induction of the browning process [5–7].

Dietary molecules promote thermogenic response primarily by activating β3-adrenergic receptors, which initiates a signaling cascade involving increased cyclic AMP and protein kinase A activation. This cascade leads to phosphorylation of downstream target molecules and stimulates lipolysis. Additionally, the activation of AMP-activated protein kinase (AMPK) and its downstream pathways, along with the coordinated actions of key transcriptional regulators as peroxisome proliferator-activated receptor-gamma coactivator-1alpha (PGC-1α), peroxisome proliferator-activated receptor gamma (PPARγ), and PR domain containing 16 (PRDM16), enhances the expression and activity of uncoupling protein 1 (UCP1). UCP1 dissipates the mitochondrial proton gradient, releasing energy as heat rather than producing adenosine triphosphate [8,9].

UCP1 is a key protein in the inner mitochondrial membrane that facilitates proton leakage, thereby generating heat. Although UCP1 serves as a hallmark of browning via UCP1-dependent thermogenesis, browning can occur via UCP1-independent pathways [10]. Notably, browning is linked to cellular redox state, and antioxidant compounds can enhance mitochondrial biogenesis and oxidative capacity in adipocytes, thereby promoting lipid oxidation and reducing obesity-associated oxidative stress. These findings expand the therapeutic potential of browning interventions by suggesting that modulation of adipose oxidative metabolism, even in the absence of increased UCP1 expression, may represent an effective strategy for improving lipid homeostasis and mitigating obesity-related complications [11–13].

Adipose tissue browning exhibits distinct characteristics across different fat deposits, reflecting the functional and metabolic heterogeneity of these tissues. BAT is highly thermogenic because of its high mitochondrial density and UCP1 expression, which are responsible for heat production. In contrast, subcutaneous depots, such as inguinal white adipose tissue (iWAT), show a greater propensity for browning than visceral depots because of their richer vascularization and denser sympathetic innervation, which facilitates thermogenic activation [14,15]. Conversely, visceral fat depots, such as epididymal white adipose tissue (eWAT), demonstrate limited browning capacity and are predominantly involved in energy storage and secretion of adipokines and pro-inflammatory cytokines [16].

Coffee, one of the most widely consumed beverages worldwide, is a rich source of bioactive compounds, such as caffeine, chlorogenic acids, and diterpenes (cafestol and kahweol), which have been associated with various metabolic effects [17,18]. Regular coffee consumption has been associated with increased lipid metabolism and adipose tissue browning, in addition to exerting antioxidant and anti-inflammatory effects [19–21]. However, despite these promising observations, a comprehensive understanding of its exact impact on adipose tissue remodeling in different white (inguinal and epididymal) and brown adipose deposits remains elusive.

This study aimed to investigate the effects of coffee consumption on thermogenic and lipid-metabolic pathways in brown, inguinal, and epididymal adipose depots, as well as their interrelationship with inflammatory and redox responses in rats.

## Materials and methods

### Preparation of coffee solution

The coffee solution was prepared following the guidelines of the Brazilian Coffee Industry Association [22] through percolation. Fifty grams of a commercial coffee powder, composed of a blend of *Coffea arabica* grains (Ponto Aralto, batch LJA088 3011 6, manufactured on 07/17/2019 at 07:11 am, Jundiaí, São Paulo, Brazil), were percolated using 500 mL of water at 90 °C, and a 100% cellulose fiber filter paper (Original, n° 12, Melitta).

The 10% coffee solution underwent freeze-drying and was stored at -80 °C until the diet food preparation. For diet formulation, an amount of 126 mL of a 10% coffee infusion (equivalent to 3.9 g of the freeze-dried sample) was added per kilogram of diet. The quantity of coffee added/kg diet was determined based on the average daily coffee consumption and total food intake of the Brazilian population, which is 163 mL (equivalent to one cup per day) and 1.290 g of food, respectively [23]. Therefore, as an adult rat consumes an average of 25 g of food per day, the daily consumption of the coffee solution for the rats was adjusted to 3.15 mL/day, equivalent to 126 mL of a 10% coffee solution per kilogram of feed. After lyophilization, 126 mL of a 10% coffee solution resulted in 3.9 g of powder. Therefore, 3.9 g of 10% lyophilized coffee solution was added per kilogram of diet. To prepare the diet, the freeze-dried coffee solution was thoroughly mixed with the other diet components until a homogeneous mixture was achieved. Subsequently, the mixture was hydrated and pelletized.

### Animals and diet

Twenty-eight male Wistar rats (Instituto de Ciências Biomédicas, USP, São Paulo, Brazil), recently weaned (21 days old), were individually housed in stainless steel cages with 12/12 light/dark cycles, at a temperature of 22 ± 1 °C. The diet was provided from 12 pm the day before to 8 am the day after, with free access to water. The experimental protocol received approval from the Animal Use Ethics Committee of the University of Brasília/UnB, Protocol No. 25/2018 on May 8, 2018, adhering to the guidelines of the National Council for the Control of Animal Experimentation (US) and the “Guide for the Care and Use of Laboratory Animals” [24].

The animals underwent a 7-day acclimatization period with the AIN-93G control diet [25]. Subsequently, the animals were divided into four experimental groups (seven rats/group) and treated with one of the described diets for approximately two months (56 days): I. Control (CT): AIN-93G diet; II. High-fat (HF): modified control diet, containing 58% lipids; III. Coffee (CT+): control diet supplemented with 126 mL of 10% freeze-dried coffee infusion solution (equivalent to 3.9 g of the freeze-dried sample)/kg of diet; IV. High-fat with coffee (HF+): high-fat diet supplemented with 126 mL of 10% lyophilized coffee infusion solution (equivalent to 3.9 g of the lyophilized sample)/kg of diet.

The lipid concentration of the high-fat diet (58%) was based on the Research Diets, Inc. Diet-Induced Obesity Model (D12492, Research Diets, Inc., New Brunswick, NJ, USA). Soybean oil (6%) and lard (52%) were used as fat sources in the high-fat diet.

At the end of the treatment period, euthanasia was performed under anesthesia with 3% isoflurane in an anesthetic chamber, followed by exsanguination through a cardiac puncture, with serum and plasma collection. Immediately after the procedure, the liver and epididymal, inguinal, and brown adipose tissue were washed with saline solution (NaCl 0.9%) at 4 °C and promptly frozen in liquid nitrogen (N₂). Subsequently, all samples were stored at -80 °C. A small section of the liver and adipose tissue was also preserved in a 4% formaldehyde solution for histological analysis.

### Diet Consumption and Weight Gain

Daily rat food consumption was determined by measuring the difference between the food supply (in grams) and the remainder, assessed using a precision scale (Shimadzu, model AUY220, Kyoto, Japan). Weight gain was assessed through weekly weighing of the animals on a digital scale (Marte, ASF11, São Paulo, SP, Brazil).

### Lipid Profile

Serum and liver concentrations of total cholesterol, high-density lipoprotein cholesterol (HDL), low-density lipoprotein cholesterol (LDL), and triglycerides were measured using commercial enzymatic-colorimetric kits (BioClin, Belo Horizonte, MG, Brazil), according to the manufacturer’s instructions. Absorbance was measured using a SpectraMax M2 microplate reader (Molecular Devices, Sunnyvale, CA, USA).

Liver lipid extraction was performed following the method described by Vieira et al. [26], with minor adaptations. Briefly, 50 mg of liver tissue was homogenized in 1 mL of isopropanol using an electric homogenizer (D1000-E, Benchmark Scientific, NJ, USA), then centrifuged at 2000 × g for 10 minutes at 4 °C.

### Hematoxylin and eosin staining and immunohistochemistry

Liver and adipose tissue samples (BAT, iWAT, and eWAT) were fixed in 10% formaldehyde. Then, the tissues were incubated for 48 h in a 10% formaldehyde and 1% sucrose buffer and were dehydrated using a concentration gradient of ethyl alcohol. The tissues were immersed in xylene and then embedded in paraffin. The tissue was sectioned to 5 μm (microtome Leica RM2125 RT, São Paulo, Brazil) and stained with Harris hematoxylin and aqueous eosin.

The cell morphology (unilocular or multilocular profile of adipocytes), adipocyte size, and fat accumulation in hepatocytes were captured using the Lab.A1 AX10 Zeizz microscopy and Axiocam ERc5s. ImageJ 1.54 version (U. S. National Institutes of Health, Bethesda, Maryland, USA) program was used with the Adiposoft 1.16 version (Centro de Pesquisa Médica Aplicada, Pamplona, Espanha) plugin to measure adipocyte area in adipose tissues.

For each histological section, three images were acquired from distinct regions to calculate the average lipid vesicle. Adipocyte counting was performed on three serial sections to ensure representative sampling. Identification of adipocyte types was conducted at 40x and 100x magnification.

The Non-Alcoholic Fatty Liver Disease (NAFLD) Score [27] was employed to characterize the NAFLD of the rat’s liver. The NAFLD Score includes the following criteria: microvesicular and macrovesicular steatosis, hypertrophy, and inflammation, adapted for rodents. The images were blindly analyzed, ensuring that the experimental groups were not disclosed during the evaluation.

A minimum of three sections of the histological images of each animal were used for liver analysis, as well as for measurements and counting of adipocytes, including both unilocular and multilocular types.

For immunohistochemistry, the presence of the UCP1 protein in inguinal adipose tissue was analyzed. The EasyLink Duo kit (EasyPath Diagnósticos, São Paulo, Brazil) was used according to the manufacturer’s instructions, with modifications. Histological sections (4 µm) were prepared on silanized, double-adhesion, positively charged slides (ImmunoSlide, EasyPath Diagnósticos, São Paulo, Brazil) and subjected to deparaffinization.

The slides were immersed in distilled water, and antigen retrieval was performed using a 10 mM sodium citrate buffer with 0.05% Tween 20 (pH 6.0) for 1 h.

The peroxidase blocker was applied for 5 min, followed by washing with Tris-Buffered Saline (TBS) and drying. The monoclonal antibody UCP1 (E9Z2V) XP Rabbit (Cell Signaling, Massachusetts, USA) was diluted at 1:50 in TBS and applied to the sections. The slides were incubated with the antibody overnight at 4 °C.

An antibody amplifier was then applied for 15 min and dried. The PolyFusion-HRP polymer was added to the sections and dried. Detection was performed using DAB for 5 min (a mixture of 1 mL DAB substrate with 1 drop of DAB chromogen), followed by washing with TBS, deionized water, and drying. Hematoxylin was used for 1 min.

The H-Score [28] was used to evaluate staining intensity based on the percentage of stained tissue in immunohistochemistry. Image analysis was performed in a blinded manner, without knowledge of the experimental groups and using at least three images of each animal.

### Determination of Transcript Levels of Proteins Related to Browning, Lipid Metabolism, and Inflammation

#### Total RNA Extraction

RNA extraction from adipose tissue was conducted using TRIzol reagent (Invitrogen, Carlsbad, CA, USA) following the manufacturer’s instructions with modifications. Homogenization was conducted in a tissue disruptor (Loccus, L-Beader 6, São Paulo, Brazil), using approximately 0.1 g of tissue per 1 mL of TRIzol reagent, in a tube previously filled with 1 mm glass or zirconium microspheres. Homogenization was made using cycles of 6 m/s for 30 s.

Following homogenization, the samples were centrifuged at 12,000 × g for 10 min at 4 °C, and the upper transparent fat layer was removed and discarded using a 200 µL tip. The red phase containing the RNA was transferred to a clean microtube and kept at room temperature for 5 min. Subsequently, 200 µL of chloroform (Merck, Darmstadt, Germany) was added to RNA, incubated for 3 min at room temperature, and then centrifuged at 10,000 × g at 4 °C for 10 min. The upper aqueous phase containing the RNA was removed, precipitated with 500 µL of isopropanol (J. T. Baker, Xalostoc, Edo de Méx, México), and centrifuged at 10,000 × g at 4 °C for 10 min. The pellet was washed with 1 mL of 75% ethanol (J. T. Baker, Xalostoc, Edo de Méx, México), centrifuged at 10,000 × g at 4 °C for 10 min, and the supernatant was discarded. The pellet was air-dried at room temperature and subsequently resuspended in 10 μL of autoclaved MilliQ water.

To enhance material quality, the samples were precipitated with sodium acetate (3 mol/L, pH 5.2, 0.1 x sample volume) and ethanol (2.5 x sample volume) for 30 min on ice. After precipitation, samples were centrifuged at 10,000 × g for 30 min at 4 °C. The supernatant was discarded, and 1 mL of 75% ethanol was added at 4 °C. Samples were centrifuged at 10,000 × g for 5 min at 4 °C, and the supernatant was discarded. The pellet was air-dried at room temperature and resuspended as previously described.

Total RNA concentration was calculated using the equation: RNA (µg/mL) = A260 nm x 40 x dilution factor. RNA sample purity was assessed using the A260 nm/A280 nm ratio (∼1.8-2.0) and A_260_/A_230_ ratio (∼2.0) [29]. RNA integrity was evaluated through electrophoretic profiling in a 0.8% agarose gel (BioAgency, São Paulo, SP, Brazil), with 1x Tris-Acetate-EDTA running buffer and stained with 5 µL GelRed^TM^ (Thermo Scientific Inc., Carlsbad, CA, USA). Electrophoretic profiles were analyzed using the L Pix HE Image photo documentation system software (Loccus Biotecnologia, SP, Brazil).

#### Quantification of mRNA by qPCR

Complementary DNA (cDNA) synthesis was conducted using the High-Capacity cDNA Reverse Transcription Kit with RNase Inhibitor (Applied Biosystems, Foster City, CA, USA). A 2X mix was prepared, comprising 10X RT Buffer, 10X Random Primers, dNTP Mix, RNase Inhibitor, and MultiScribe^TM^ Reverse Transcriptase. Subsequently, 10 µL of the 2X mix was added to a microtube containing 10 µL of the RNA sample at a concentration of 0.2 µg/µL. Reverse transcription was performed using a thermocycler (Esco, Swift Maxi, Hatboro, PA, USA) with the following parameters: 25 °C for 10 min; 37 °C for 120 min; 85 °C for 5 min. After the reaction, cDNA samples were stored at -20°C. A cDNA synthesis reaction without the reverse transcriptase enzyme was performed for each sample as a negative control. Subsequently, an aliquot of the sample without reverse transcriptase was subjected to a real-time polymerase chain reaction (qPCR) to confirm the absence of genomic DNA contamination.

The qPCR reaction was composed of 2 µL of cDNA, 5 µL of SYBR Mix, and 0.2 µmol/L (final concentration) of each primer (Table 1) for a final volume of 10 µL. The amplification program was performed for PowerTrack SYBR Green Master Mix: 1 reaction cycle at 95° for 2 min (enzymatic activation), followed by 40 reaction cycles (Denaturation: 95 °C for 5 s; Annealing and extension: 60 °C for 30 s) and dissociation curve stage (Melting Curve): 95 °C for 15 s, 60 °C for 60 s, 95 °C for 15 s. For the SYBR Green PCR Master Mix, 1 reaction cycle was used at 95 °C for 10 min (enzymatic activation), followed by 40 reaction cycles (Denaturation: 95 °C for 15 s; Annealing and extension: 60 °C for 60 s) and dissociation curve stage (melting curve): 95 °C for 15 s, 60 °C for 60 s, 95 °C for 15 s. The dosages were conducted in duplicate, and the specificity of each amplified product was verified using the dissociation curve (melting curve). The 2-ΔΔCT method [30] was used to calculate the relative expression of each gene, with the estimate of the ΔΔCT value being based on the difference between the ΔCt value (amplification cycle of the gene of interest - cycle amplification of the constitutive β-actin gene) of the test group and the ΔCt of the control group.

**Table 1.**
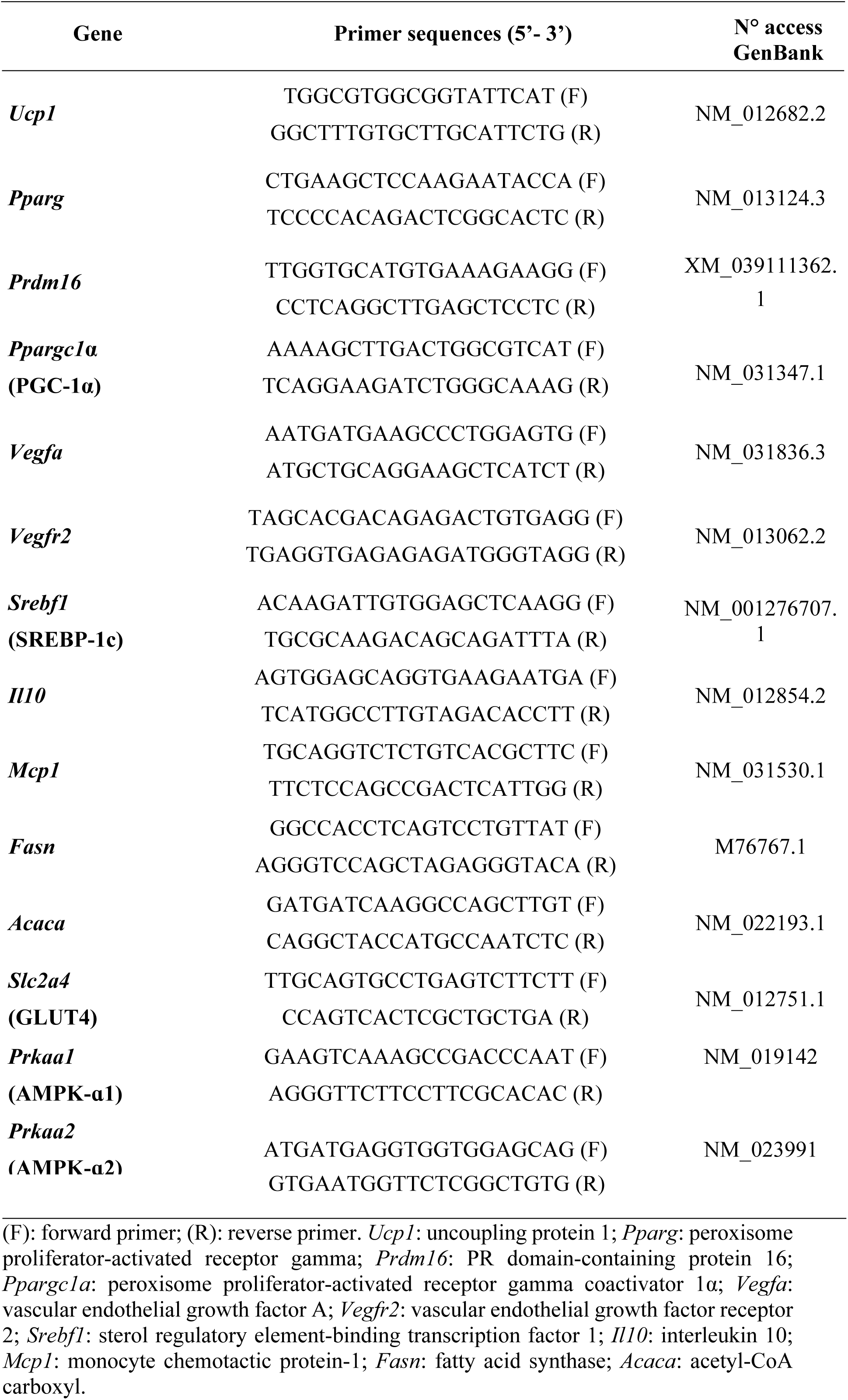
Sequences of primers used for qPCR assay.

### Measurement of Oxidative Damage and Antioxidant Capacity

#### Carbonylated Proteins

The concentration of carbonylated proteins in BAT, iWAT, and eWAT was determined by detecting the complex formed between carbonyl groups of cellular proteins and the reagent 2,4-dinitrophenylhydrazine (DNPH) [28] at a wavelength of 450 nm using a UV-1800 UV-VIS spectrophotometer (Shimadzu, Kyoto, Japan). The concentration of carbonylated proteins was determined using the molar extinction coefficient of the protein-DNPH complex, which is 22.308 mmol/L^-1^cm^-1^ at 450 nm. The total protein concentration of the homogenates was determined using the Hartree method [31], and the results were expressed as nmol of carbonyl/mg of total protein.

#### Lipid Peroxidation

The level of lipid peroxidation in adipose tissue was determined using the method described by Candan and Tuzmen [32]. The technique relies on the reaction of malonaldehyde (MDA) with thiobarbituric acid. This complex is detected using a wavelength of 553 nm for excitation and 532 nm for emission monitoring in a spectrofluorometer (Spectramax M2, USA). MDA concentration was determined using a standard curve constructed from the hydrolysis of the 1,1,3,3-tetraethoxy-propane 97% (TEP; Sigma Aldrich, St Louis, MO, USA) in 1% sulfuric acid at concentrations of 0; 0.3125; 0.6250; 1.2625; 2,525; 5.05 nmol/mL. The MDA concentration was expressed as nmol of MDA/mg tissue.

#### Specific Activity of Antioxidant Enzymes

The activities of the enzymes catalase (CAT), glutathione peroxidase (GPx), glutathione-S-transferase (GST), and superoxide dismutase (SOD) were determined in the animals’ adipose tissue. The tissue was homogenized in 50 mmol/L potassium phosphate buffer (pH 7.2), containing 0.5 mmol/L ethylenediamine tetraacetic acid (EDTA) and 1 mmol/L phenylmethylsulfonyl fluoride (PMSF), in a tissue x solution ratio of 1:7.5 (p/v), using a handheld homogenizer TissueRuptor (QIAGEN, Austin, Texas, USA), and samples were homogenized on ice. The homogenate was centrifuged at 10,000 × g at 4 °C for 20 min, and the supernatant was used to determine enzymatic activities and total protein.

The specific activity of the CAT enzyme was determined according to the method described by Joanisse and Storey [33]. The reaction system consisted of 30 μL of adipose tissue homogenate, 50 mmol/L potassium phosphate buffer (pH 7.2), 0.5 mmol/L EDTA, and 10 mmol/L H_2_O_2_, for a final reaction volume of 500 μL completed with H_2_O MilliQ. H_2_O_2_ consumption will be monitored at 240 nm/30 s. One unit of CAT is defined as the number of enzymes required to decompose 1 μmol of H_2_O_2_/min.

The specific activity of the GPx enzyme was determined by monitoring the oxidation of nicotinamide adenine dinucleotide phosphate (NADPH) at 340 nm/30 s [33]. The reaction system with a final volume of 500 μL was composed of potassium phosphate buffer 50 mmol/L (pH 7.2), EDTA 0.5 mmol/L, NaN_3_ 2 mmol/L, glutathione reductase 1.5 IU/mL, NADPH 0.15 mmol/L, reduced glutathione 5 mmol/L, H_2_O_2_ 0.2 mmol/L final concentrations, and 30 μL of adipose tissue homogenate, with the final volume completed with H_2_O MilliQ. The unit (U) of GPx is defined as the number of enzymes required to oxidize 1 nmol NADPH/min.

GST activity was quantified at 340 nm/30 s by monitoring the formation of the 2,4-dinitrophenyl-glutathione complex. The final volume reaction of 500 µL is composed of 50 mmol/L potassium phosphate buffer (pH 7.2), 0.5 mmol/L EDTA, 1 mmol/L 1-chloro-2,4-dinitrobenzene, 1 mmol/L reduced glutathione (final concentration values), and 50 µL of adipose tissue homogenate. One unit (U) of GST is defined as the number of enzymes required to produce 1 nmol of product/min [34].

### Statistical Analysis

Values were presented as least squares means ± confidence interval limits (n = 4-7). The effects of diets (control and high-fat), coffee (with or without), and the diet x coffee interaction were analyzed in a 2 × 2 factorial design. Homogeneity of variances between treatments was assumed. Significant interactions between diet and coffee were further explored and compared using Tukey’s test. Outliers were removed using the box plot method. Statistical analyses were performed with the PROC GENMOD procedure from SAS/STAT® (SAS OnDemand, SAS Institute Inc., Cary, NC, USA), and statistical significance was set at p < 0.05.

## Results

### Physiological variables and lipid profile

Diet effects were observed for food consumption and body weight gain (p < 0.001; Table 2), and rats fed the high-fat (HF) diet showed higher body weight gain and lower food intake than those of the control (CT) rats. Coffee consumption had no significant effect.

**Table 2.** Physiological variables of rats treated with control (CT) or high-fat diet (HF) with (+) or without (-) coffee.

| Diet | Coffee |  | Mean (95% CI) | Two-way ANOVA P values |  |  |
| --- | --- | --- | --- | --- | --- | --- |
|  | (-) | (+) |  | Diet | Coffee | Diet x Coffee |
| Total food consumption (g/56 day) |  |  |  |  |  |  |
| CT | 1,021.7 (962.1 - 1,081.3) | 1,062.2 (1,002.7 - 1,121.8) | 1,042.0 (999.8 - 1,084.1) <sup>a</sup> | <0.001 | 0.748 | 0.106 |
| HF | 912.1 (852.5 - 971.6) | 852.0 (792.4 - 911.6) | 882.0 (839.9 - 924.1) <sup>b</sup> |  |  |  |
| Mean | 966.9 (924.8 - 1009.0) | 957.1 (915.0 - 999.2) |  |  |  |  |
| Body weight gain (g) |  |  |  |  |  |  |
| CT | 268.9 (242.2 - 295.6) | 273.1 (246.4 - 299.9) | 271.0 (252.1 - 289.9) <sup>b</sup> | <0.001 | 0.596 | 0.402 |
| HF | 342.6 (315.9 - 369.3) | 323.9 (297.2 - 350.6) | 333.2 (314.4 - 352.1) <sup>a</sup> |  |  |  |
| Mean | 305.8 (286.9 - 324.7) | 298.5 (279.6 - 317.4) |  |  |  |  |

Values are expressed as mean ± confidence limits. Means differ ( $P < 0.05$ ); $n = 7$ . CT: rats fed with control diet (AIN-93G); HF: rats fed with high-fat diet; (+): rats fed with coffee; (-): rats fed without coffee. Capital letters indicate differences between values in the same row, while lowercase letters indicate differences in the same column.

Regarding the serum lipid profile (Table 3), the HF diet reduced the serum HDL concentration and LDL/HDL ratio (p = 0.026) compared to those with the CT diet. The HF+ diet promoted an increase in the serum LDL/HDL ratio compared to that with the HF diet (p < 0.0001).

**Table 3.** Lipid profile of rats treated with control (CT) or high-fat diet (HF) with (+) or without (-) coffee.

| Diet | Coffee |  | Mean (95% CI) | Two-way ANOVA P values |  |  |
| --- | --- | --- | --- | --- | --- | --- |
|  | (-) | (+) |  | Diet | Coffee | Diet x Coffee |
| Serum lipid profile |  |  |  |  |  |  |
| Total cholesterol |  |  |  |  |  |  |
| CT | 70.4 (54.5 - 86.3) | 78.0 (63.5 - 92.5) | 74.2 (63.4 - 85.0) | 0.662 | 0.975 | 0.351 |
| HF | 74.3 (58.4 - 90.3) | 67.3 (52.7 - 81.8) | 70.8 (60.0-81.6) |  |  |  |
| Mean | 72.4 (61.1 - 83.6) | 72.6 (62.4 - 82.9) |  |  |  |  |
| Triglycerides |  |  |  |  |  |  |
| CT | 105.8 (83.4 - 128.2) | 95.5 (62.1 - 122.9) | 100.7 (83.0 - 118.3) | 0.968 | 0.241 | 0.743 |
| HF | 108.7 (86.4 - 131.1) | 90.7 (70.0 - 111.4) | 99.7 (84.5 - 115.0) |  |  |  |
| Mean | 107.3 (91.5 - 123.1) | 93.1 (75.9 - 110.3) |  |  |  |  |
| Low-density lipoprotein |  |  |  |  |  |  |
| CT | 22.4 (16.8 - 27.9) | 20.3 (14.7 - 25.8) | 21.3 (17.4 - 25.2) | 0.459 | 0.562 | 0.198 |
| HF | 16.5 (11.0 - 22.1) | 21.9 (16.3 - 27.4) | 19.2 (15.3 - 23.1) |  |  |  |
| Mean | 19.4 (15.5 - 23.4) | 21.1 (17.2 - 25.0) |  |  |  |  |
| High-density lipoprotein |  |  |  |  |  |  |
| CT | 69.0 (57.3 - 80.7) | 78.8 (67.1 - 90.5) | 73.9 (65.6 - 82.1) <sup>a</sup> | 0.026 | 0.779 | 0.070 |
| HF | 66.1 (54.4 - 77.8) | 52.9 (40.1 - 65.7) | 59.5 (50.9 - 68.2) <sup>b</sup> |  |  |  |
| Mean | 67.6 (59.3 - 75.8) | 65.8 (57.2 - 74.5) |  |  |  |  |
| LDL/HDL ratio |  |  |  |  |  |  |
| CT | 0.34 (0.28 - 0.38) <sup>a</sup> | 0.25 (0.20 - 0.30) | 0.29 (0.26 - 0.33) | 0.040 | 0.057 | < 0.001 |
| HF | 0.25 (0.20 - 0.30) <sup>B,b</sup> | 0.45 (0.39 - 0.51) <sup>A</sup> | 0.31 (0.31 - 0.39) |  |  |  |
| Mean | 0.29 (0.26 - 0.33) | 0.35 (0.31 - 0.39) |  |  |  |  |
| Hepatic lipid profile |  |  |  |  |  |  |
| Total cholesterol |  |  |  |  |  |  |
| CT | 8.5 (3.2 - 13.7) | 14.7 (10.2 - 19.1) | 11.6 (8.1 - 15.0) <sup>b</sup> | < 0.001 | 0.015 | 0.994 |
| HF | 19.8 (15.0 - 24.6) | 26.1 (21.6 - 30.5) | 22.9 (19.7-26.2) <sup>a</sup> |  |  |  |
| Mean | 14.1 (10.6 - 17.7) <sup>B</sup> | 20.4 (17.2 - 23.5) <sup>A</sup> |  |  |  |  |
| Triglycerides |  |  |  |  |  |  |
| CT | 77.2 (51.1 – 103.3) | 127.8 (98.6 – 157.0) | 102.5 (82.9 – 122.1) <sup>b</sup> | < 0.001 | 0.015 | 0.357 |
| **HF** | 155.8 (129.7 - 181.9) | 180.1 (150.9 – 209.3) | 167.9 (148.4 - 187.5) <sup>a</sup> |  |  |  |
| **Mean** | 116.5 (98.1 - 135.0) <sup>B</sup> | 154.0 (133.3 – 174.6) <sup>A</sup> |  |  |  |  |
| **Low-density lipoprotein** |  |  |  |  |  |  |
| **CT** | 4.5 (3.4 - 5.5) | 5.2 (4.3 - 6.2) | 4.6 (3.9 - 5.3) | 0.564 | 0.752 | 0.237 |
| **HF** | 4.8 (3.9 - 5.7) | 4.4 (3.3 - 5.4) | 4.8 (4.1 - 5.5) |  |  |  |
| **Mean** | 4.9 (4.2 - 5.6) | 4.6 (3.9 - 5.3) |  |  |  |  |
| **High-density lipoprotein** |  |  |  |  |  |  |
| **CT** | 7.5 (6.7 - 8.4) | 7.9 (7.1 - 8.7) | 7.7 (7.2 - 8.3) | 0.332 | 0.510 | 0.819 |
| **HF** | 8.0 (7.2 - 8.9) | 8.0 (7.2 - 8.9) | 8.1 (7.5 - 8.7) |  |  |  |
| **Mean** | 7.8 (7.2 - 8.4) | 8.1 (7.5 - 8.6) |  |  |  |  |
| **LDL/HDL ratio** |  |  |  |  |  |  |
| **CT** | 0.62 (0.47 - 0.78) | 0.68 (0.54 - 0.82) | 0.62 (0.51 - 0.73) | 0.341 | 0.847 | 0.381 |
| **HF** | 0.62 (0.47 - 0.77) | 0.54 (0.38 - 0.69) | 0.61 (0.50 - 0.71) |  |  |  |
| **Mean** | 0.65 (0.55 - 0.75) | 0.58 (0.47 - 0.68) |  |  |  |  |

Values are expressed as mean ± confidence limits. Means differ ( $P < 0.05$ ); n= 4-7/group. CT: rats fed with control diet (AIN-93G); HF: rats fed with high-fat diet; (+): rats fed with coffee; (-): rats fed without coffee; HDL: high-density lipoprotein; LDL: low-density lipoprotein. Capital letters indicate differences between values in the same row, while lowercase letters indicate differences in the same column.

In the liver, the HF diet consumption promoted an increase in total cholesterol (p < 0.001) and triglycerides (p = 0.001) compared to those with the CT diet. Coffee exhibited an effect on total cholesterol and triglycerides in the liver (p = 0.015 for both), with coffee consumption promoting an increase in both levels regardless of diet type.

### Area of adipocytes

Fig 1 shows hematoxylin and eosin-stained microscopic images of adipose tissues and livers of rats fed a CT or HF diet, with (+) or without (-) coffee. In BAT and iWAT, a diet with coffee interaction was observed, with HF diet-fed rats showing a larger adipocyte area than those fed the CT diet (1,002.64 and 509.24 μm², p = 0.0037; 4,482.87 and 2,557.20 μm², p < 0.0001, respectively; Fig 1). The adipocyte area of rats fed the HF+ diet was smaller than those fed the HF diet (372.87 and 509.24 μm²; 2,608.62 and 4,482,87, respectively; p < 0.0001).

**Fig 1.**
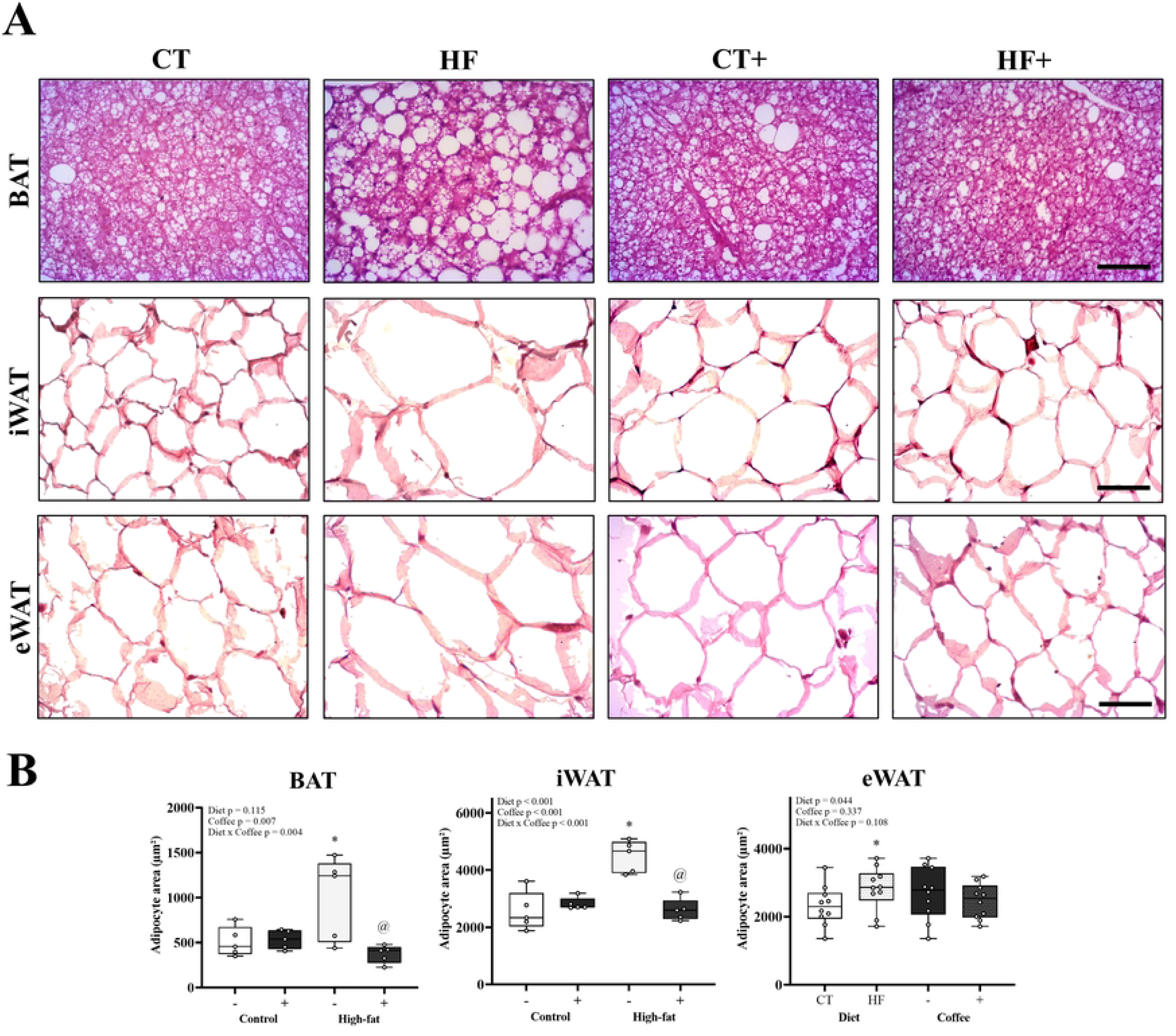
Histopathological analysis of adipose tissue from rats. (A) Morphological evaluation of brown (BAT), inguinal (iWAT), and epididymal (eWAT) adipose tissues from rats fed a control (CT), high-fat (HF) diets, with (+) or without (-) coffee. Stain: hematoxylin and eosin. Scale bar: 20 µm in BAT; 10 µm in eWAT and iWAT. (B) Quantification of adipocyte area of BAT, iWAT, and eWAT. Values are mean ± confidence limits (n = 5/group). If there is no Diet x Coffee interaction, the x-axis: Diet (CT or HF) and Coffee (-) or (+); when present, Control (-) or (+) and High-fat (-) or (+). Statistical symbols: * Control *versus* high-fat (main effect of diet); # With coffee *versus* without coffee (main effect of coffee). For interactions: * CT (-) *vs.* HF (-); & CT (-) *vs.* CT (+); @ HF (-) vs. HF (+); P < 0.05.

In eWAT, only a diet effect (p = 0.044) was observed, with the HF diet-fed rats showing a large adipocyte area than those treated with a CT diet (2,825.87 and 2,321.68 μm², respectively; Fig 1).

### Multilocular and unilocular adipocyte counting and immunohistochemistry of iWAT

In BAT, a diet with coffee interaction was observed in the number of unilocular adipocyte. The HF diet increased the number of unilocular adipocytes compared to that with the CT diet (90.99 and 34.33, respectively; p < 0.0001, Fig 2), whereas the HF+ diet promoted a decrease in the number of multilocular adipocytes compared to that with the HF diet (p < 0.0001).

**Fig 2.**
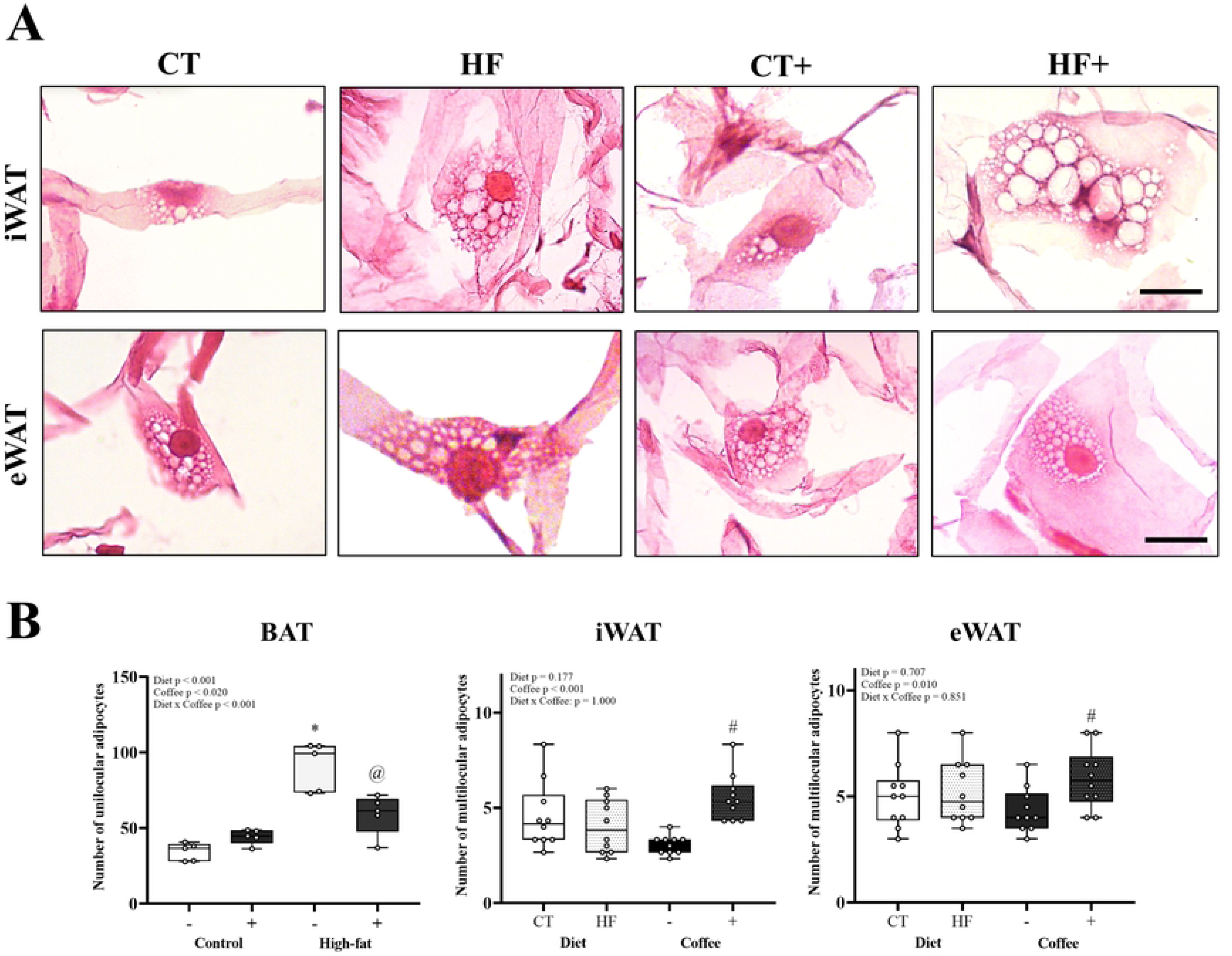
Multilocular and unilocular adipocytes. (A) Multilocular adipocytes in inguinal (iWAT) and epididymal (eWAT) adipose tissues. Stain: hematoxylin and eosin. Scale bar: 5 µm; (B) Quantification of unilocular adipocytes in brow adipose tissue (BAT) and multilocular adipocytes in iWAT and eWAT from rats fed a control (CT), high-fat (HF) diets, with (+) or without (-) coffee. Values are mean ± confidence limits (n = 5/group). If there is no Diet x Coffee interaction, the x-axis: Diet (CT or HF) and Coffee (-) or (+); when present, Control (-) or (+) and High-fat (-) or (+). Statistical symbols: * Control *versus* high-fat (main effect of diet); # With coffee *versus* without coffee (main effect of coffee). For interactions: * CT (-) *vs.* HF (-); & CT (-) *vs.* CT (+); @ HF (-) vs. HF (+); P < 0.05.

Coffee consumption affected the number of multilocular adipocytes in the iWATs and eWATs, with coffee consumption promoting an increase in the number of multilocular adipocytes (3.06 and 5.53; p < 0.0001 and 4.35 and 5.85; p = 0.010, respectively; Fig 2), regardless of diet type.

In the H-score analysis of anti-UCP1 immunostaining intensity in iWAT images (Fig 3), coffee exhibited an effect (p = 0.038), with coffee consumption promoting an increase in the score regardless of diet type.

**Fig 3.**
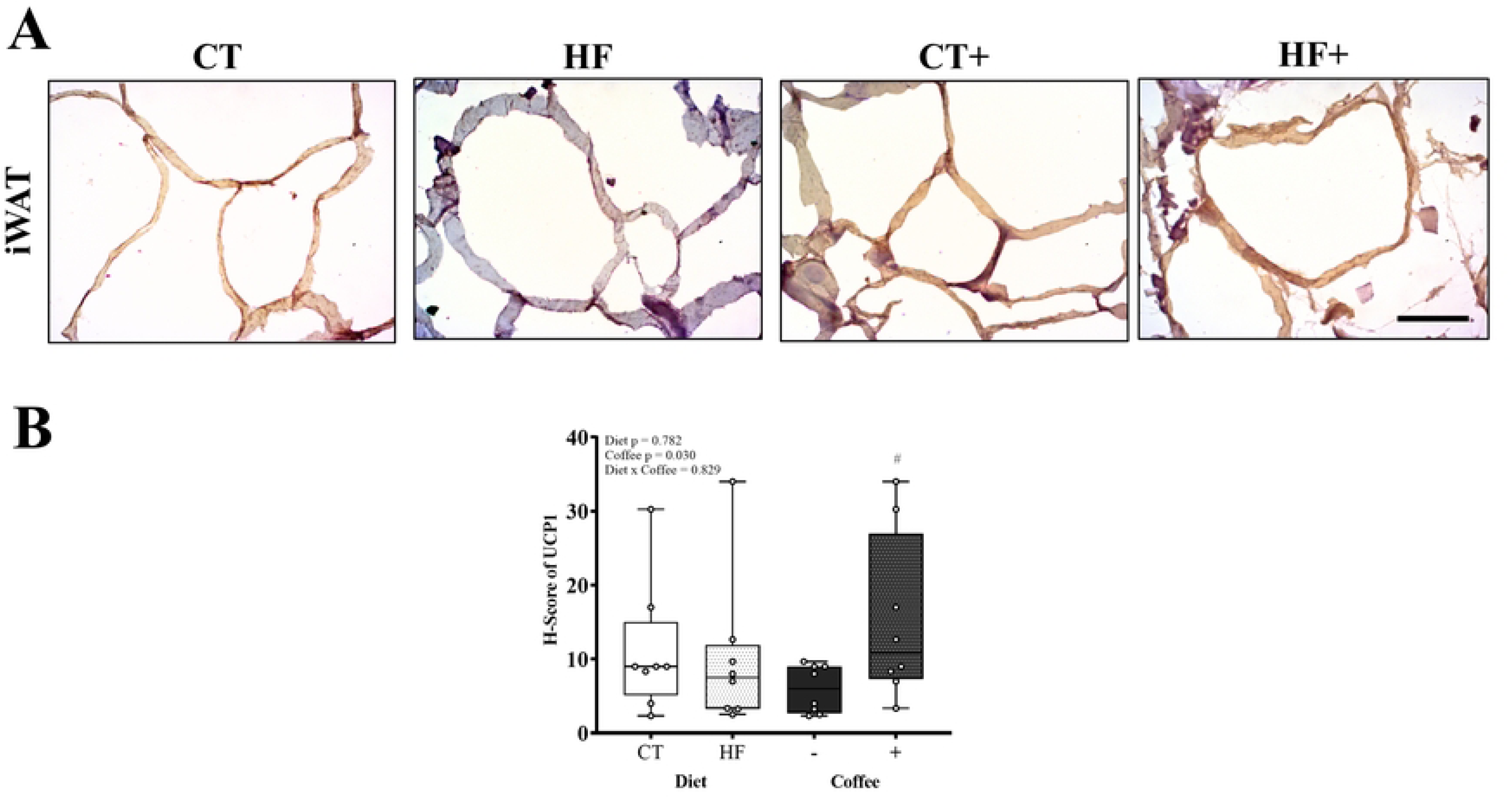
UCP1 immunostaining in inguinal adipose tissue. (A) Adipocyte immunostained with UCP1 in iWAT. Counterstain: hematoxylin. Scale bar = 5 µm. (B) Histoscore (H-score) quantification of UCP1 immunostaining in iWAT from rats fed a control (CT), high-fat diet (HF), without (-) or with (+) coffee. Values are mean ± confidence limits (n = 4/group). Statistical symbols: * Control *versus* high-fat (main effect of diet); # With coffee *versus* without coffee (main effect of coffee).

### Liver steatosis score

Fig 4 shows the scores for each criterion used to characterize the NAFLD in rats fed a CT or HF diet, with (+) or without (-) coffee. Only the HF group scored macrovesicles, with no statistical differences between the groups. A diet x coffee effect was observed on the hepatic microvesicular score (p < 0.001), with the CT+ diet promoting an increase in the microvesicular score compared to that with the CT diet (p < 0.001). The HF diet consumption also increased the microvesicular score compared to that with the CT diet (p < 0.0001).

**Fig 4.**
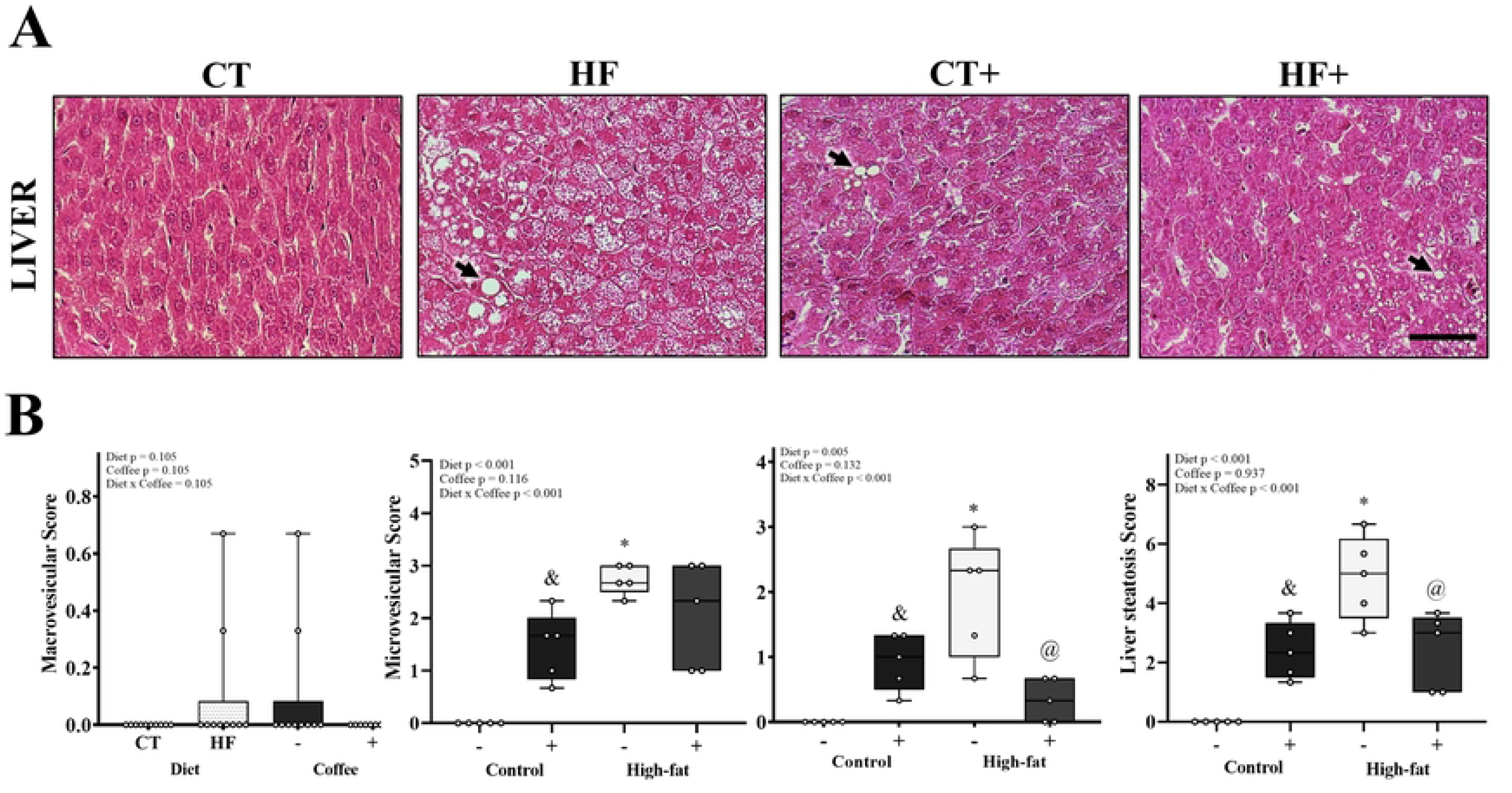
Liver histology and steatosis scoring. (A) Histological sections of liver tissue. Arrows indicate microvesicular steatosis. Stain: hematoxylin and eosin. Scale bar: 10 μm. (B) Quantification of macrovesicular steatosis, microvesicular steatosis, hepatocyte hypertrophy, and liver steatosis score from rats fed a control (CT), high-fat diet (HF), with (+) or without (-) coffee. Values are mean ± confidence limits (n = 5/group). If there is no Diet x Coffee interaction, the x-axis: Diet (CT or HF) and Coffee (-) or (+); when present, Control (-) or (+) and High-fat (-) or (+). Statistical symbols: * Control *versus* high-fat (main effect of diet); # With coffee *versus* without coffee (main effect of coffee). For interactions: * CT (-) *vs.* HF (-); & CT (-) *vs.* CT (+); @ HF (-) vs. HF (+); P < 0.05.

A diet x coffee interaction was obtained for Hypertrophy Score and Liver Steatosis Score, which showed an increase in CT+ group relative to that with the CT group (p = 0.002 and 0.001), and the HF group compared to that of the CT (p < 0.0001 for both), and a reduction in the HF+ group relative to the HF group (p < 0.0001 for both).

### mRNA levels of genes involved in lipid metabolism in BAT, iWAT, and eWAT

The mRNA levels of genes related to lipid metabolism are shown in Fig 5. In BAT, there was an effect of diet, and coffee consumption regardless of the diet type, the HF diet and coffee consumption promoted a decrease in the mRNA levels of *Acaca* (p = 0.001 and p = 0.001, respectively) and *Fasn* (p < 0.0001 and p = 0.020, respectively) compared to that with the CT diet and to that with no coffee, respectively. There were no changes in the mRNA levels of *Slc2a4*, *Prkaa1,* or *Prkaa2* after any treatment.

**Fig 5.**
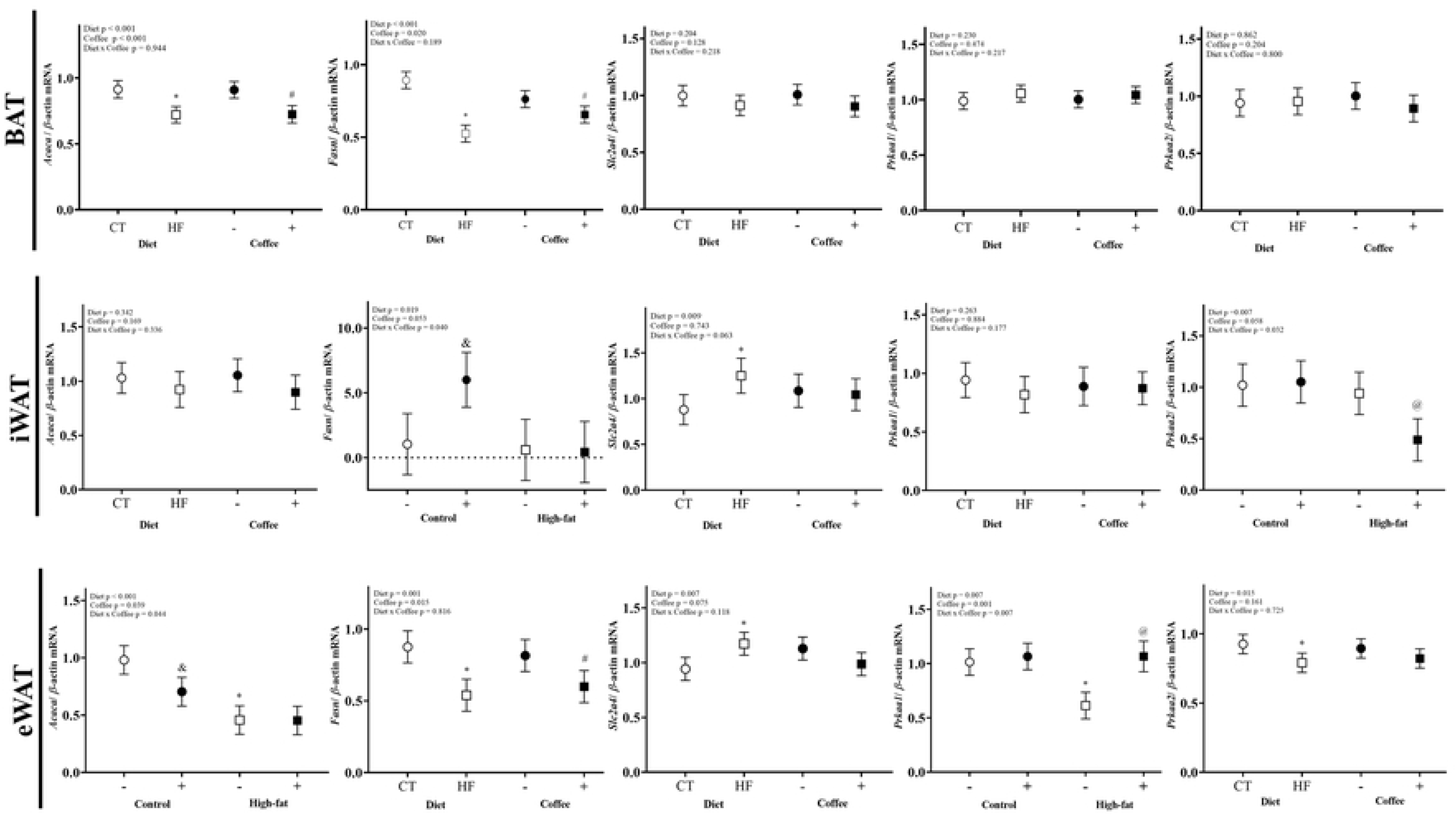
mRNA levels of genes in BAT, iWAT, and eWAT from rats. Values are mean ± confidence limits (n = 4-7/group). Control (CT), high-fat diet (HF), with (+) or without (-) coffee. If there is no Diet x Coffee interaction, the x-axis: Diet (CT or HF) and Coffee (-) or (+); when present, Control (-) or (+) and High-fat (-) or (+). Statistical symbols: * Control *versus* high-fat (main effect of diet); # With coffee *versus* without coffee (main effect of coffee). For interactions: * CT (-) *vs.* HF (-); & CT (-) *vs.* CT (+); @ HF (-) vs. HF (+); P < 0.05.

In the iWAT, there was a six-fold increase in *Fasn* mRNA levels due to coffee consumption compared to that with no coffee (p = 0.011). The HF diet increased the *Slc2a4* mRNA levels relative to that with the CT diet (p = 0.009). The diet x coffee interaction had an effect on *Prkaa2* mRNA levels (p = 0.032); the HF+ diet promoted a reduction compared to that with the HF diet (p = 0.012). The mRNA levels of *Acaca* and *Prkaa1* were not significantly different between groups.

In the eWAT, a diet x coffee interaction was obtained in mRNA levels of *Acaca* and *Prkaa1* (p = 0.044 and 0.007, respectively). *Acaca* mRNA levels was reduced in the CT+ diet compared to that with the CT diet (p = 0.011). The HF diet also decreased *Acaca* mRNA levels compared to that with the CT diet (p = 0.027). The HF+ diet increased its *Prkaa1* mRNA levels compared to that with the HF diet (p < 0.0001), whereas the HF reduced its levels compared to that with the CT diet (p < 0.0001). A dietary effect was observed on *Slc2a4* and *Prkaa2* mRNA levels (p = 0.007 and p = 0.015, respectively). The HF diet increased *Slc2a4* mRNA levels and decreased *Prkaa2* mRNA levels compared to that with the CT diet (p = 0.007 and 0.015, respectively). *Fasn* mRNA levels reduced with the HF diet compared to that with the CT diet, and with coffee consumption compared to that with no coffee (p = 0.001 and 0.015, respectively).

### mRNA levels of thermogenesis-related genes in BAT, iWAT, and eWAT

Fig 6 shows the mRNA levels of the thermogenesis-related genes in different adipose tissues. In BAT, coffee consumption reduced *Pparg* mRNA levels (p = 0.018), regardless of the diet type. Diet had an effect on *Ppargc1a* and *Ucp1* genes, with the HF diet promoting an increase in *Ppargc1a* and *Ucp1* mRNA levels compared to that with the CT diet (p = 0.001 and p < 0.0001, respectively). Neither the type of diet nor coffee consumption altered *Srebf1* and *Prdm16* mRNA levels in BAT.

**Fig 6.**
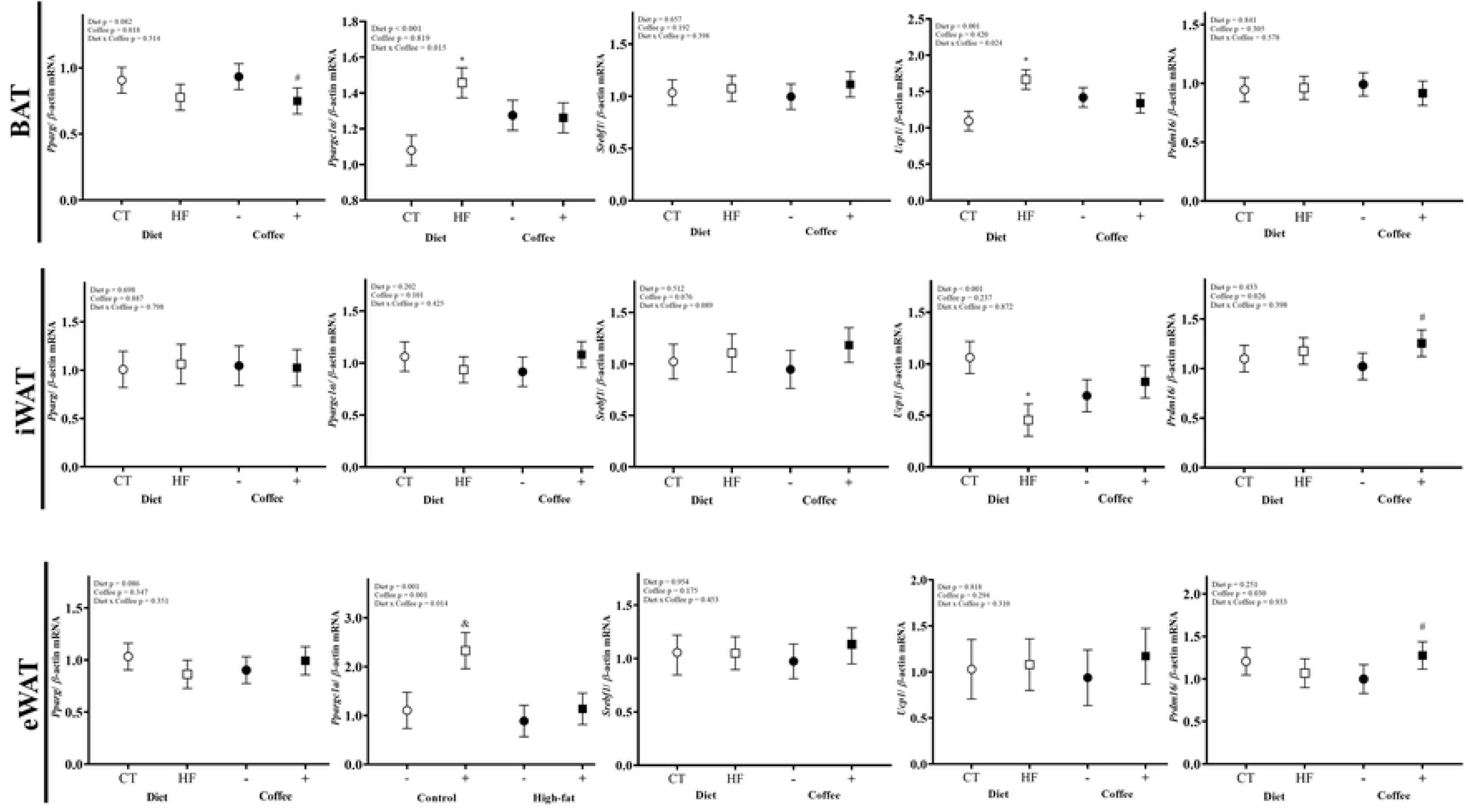
mRNA levels of thermogenic genes in BAT, iWAT, and eWAT from rats. Values are mean ± confidence limits (n = 4-7/group). Control (CT), high-fat diet (HF), with (+) or without (-) coffee. If there is no Diet x Coffee interaction, the x-axis: Diet (CT or HF) and Coffee (-) or (+); when present, Control (-) or (+) and High-fat (-) or (+). Statistical symbols: * Control *vs.* high-fat (main effect of diet); # With coffee *vs.* without coffee (main effect of coffee). For interactions: * CT (-) *vs.* HF (-); & CT (-) *vs.* CT (+); @ HF (-) vs. HF (+); P < 0.05.

In iWAT, the effect of diet type on *Ucp1* was observed (p < 0.001), with a HF diet reducing *Ucp1* mRNA levels by 57.1% compared to that with the CT diet. Regarding *Prdm16*, coffee consumption promoted an increase (p = 0.026) in *Prdm16* mRNA levels regardless of diet. No differences were found for *Pparg*, *Ppargc1a,* and *Srebf1* mRNA levels.

Regarding thermogenic genes in eWAT, an effect of the diet x coffee interaction was observed in *Ppargc1a* (p = 0.014), with the CT+ diet promoting an increase in *Ppargc1a* mRNA levels compared to that with the CT diet (p < 0.0001). Coffee consumption promoted an increase in *Prdm16* mRNA levels (p = 0.030) compared to that with no coffee. No differences were obtained for *Pparg*, *Srebf1,* and *Ucp1* mRNA levels.

### mRNA levels of angiogenesis-related genes in BAT, iWAT, and eWAT

Regarding angiogenesis-related genes (Fig 7) in BAT, the HF diet increased the mRNA levels of *Vegfa* and *Vegfr2* compared to that with the CT diet (p = 0.008 and 0.001, respectively). Coffee consumption also increased *Vegfr2* mRNA levels compared to that with no coffee (p = 0.009).

**Fig 7.**
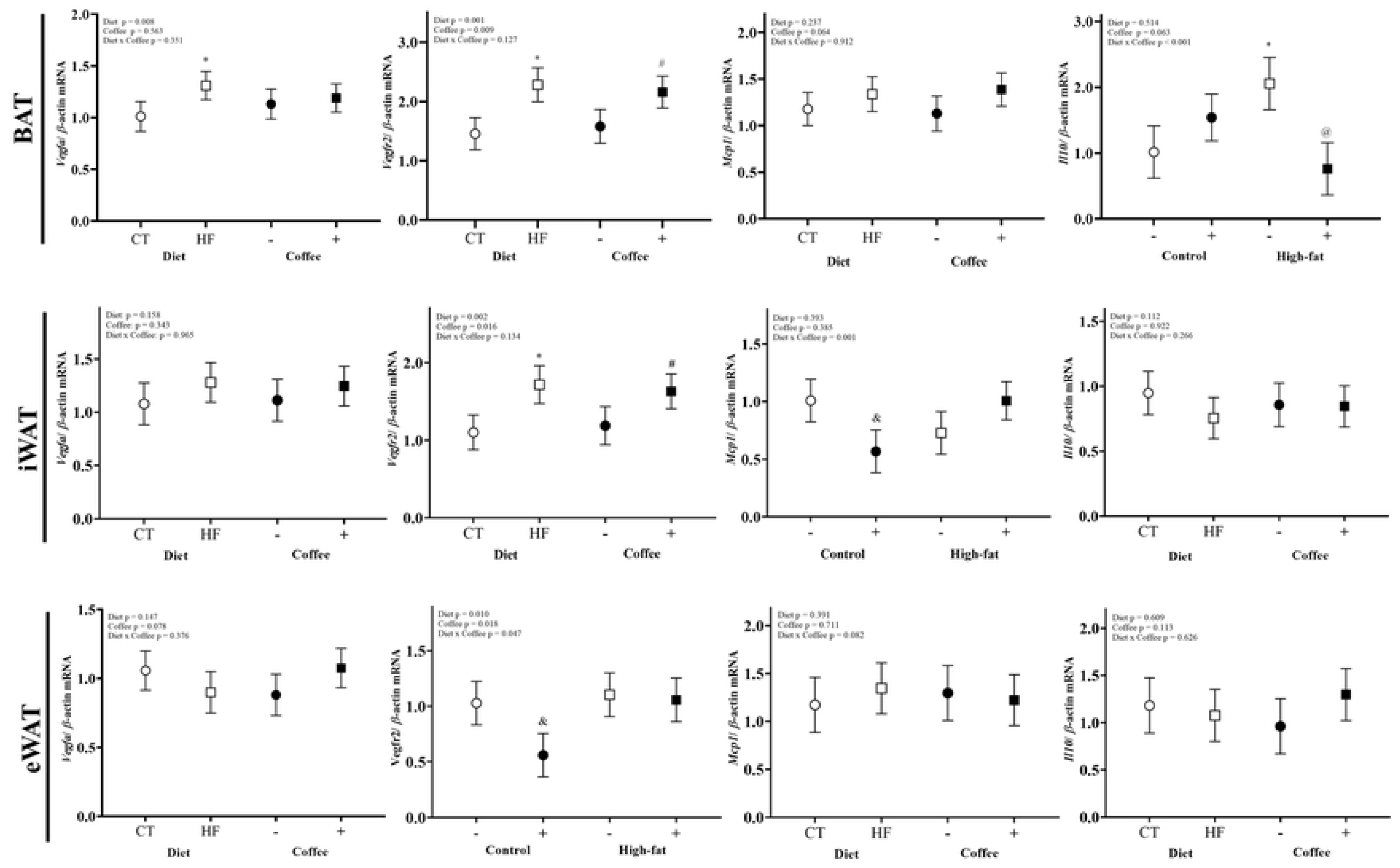
mRNA levels of angiogenesis and inflammatory genes in BAT, iWAT, and eWAT from rats. Values are mean ± confidence limits (n = 4-7/group). Control (CT), high-fat diet (HF), with (+) or without (-) coffee. If there is no Diet x Coffee interaction, the x-axis: Diet (CT or HF) and Coffee (-) or (+); when present, Control (-) or (+) and High-fat (-) or (+). Statistical symbols: * Control *vs.* High-fat (main effect of diet); # With coffee *vs.* without coffee (main effect of coffee). For interactions: * CT (-) *vs.* HF (-); & CT (-) *vs.* CT (+); @ HF (-) vs. HF (+); P < 0.05.

A similar profile was observed in the iWAT; the HF diet and the coffee consumption regardless of the diet type increased *Vegfr2* mRNA levels compared to that with the CT diet (p = 0.002) and to that with no coffee (p = 0.016), respectively. However, no diet type or coffee consumption effect was observed for *Vegfa* mRNA levels.

In the eWAT, diet x coffee exhibited an effect in the *Vegfr2* gene (p = 0.047), with the CT+ diet decreasing *Vegfr2* mRNA levels compared to that with the CT diet (p = 0.005). No differences in *Vegfa* mRNA levels were observed according to diet or coffee consumption.

### mRNA levels of inflammatory-related genes in BAT, iWAT, and eWAT

Concerning the inflammatory response (Fig 7) in BAT, coffee consumption marginally increased *Mcp1* mRNA levels (p = 0.064), regardless of the diet type. A diet x coffee effect was obtained on *Il10* (p < 0.001), with the HF+ diet reducing *Il10* mRNA levels compared to that with a HF diet (p < 0.0001). The HF diet upregulated *Il10* mRNA levels compared to that with the CT diet (p = 0.002).

In the iWAT, a diet x coffee effect was observed on *Mcp1* gene (p = 0.001), the CT+ diet reduced *Mcp1* mRNA levels when compared to the CT diet (p = 0.005). No significant differences were observed between the treatments.

No effect of diet type or coffee consumption was observed on *Mcp1* and *Il10* mRNA levels in eWAT.

### Oxidative damage to proteins and lipids

The oxidative damage results are presented in Table 4. A diet x coffee interaction was obtained on BAT (p < 0.001), with the HF+ diet promoting a marked increase in protein carbonyl levels relative to that with the HF diet (p < 0.0001), whereas the HF diet reduced protein carbonyl levels compared to that with the CT diet (p < 0.0001). A similar profile was observed in the eWAT; the HF+ diet increased protein carbonyl levels compared to that with the HF diet (p < 0.0001). In contrast to in BAT, the HF diet increased protein carbonyl levels in iWAT when compared to that in the CT diet.

**Table 4.** Oxidative damage to proteins and lipids in adipose tissues of rats treated with control (CT) or high-fat diet (HF) with (+) or without (-) coffee.

| Diet | Coffee |  | Mean (95% CI) | Two-way ANOVA P values |  |  |
| --- | --- | --- | --- | --- | --- | --- |
|  | (-) | (+) |  | Diet | Coffee | Diet x Coffee |
| Carbonyl (nmol/ mg ptn) |  |  |  |  |  |  |
| Brown |  |  |  |  |  |  |
| CT | 11.7 (10.1 - 13.4) <sup>a</sup> | 10.6 (8.7 - 12.4) | 11.2 (9.9 - 12.4) | 0.055 | 0.001 | <0.001 |
| HF | 5.1 (3.2 - 6.9) <sup>B, b</sup> | 13.5 (11.7 - 15.4) <sup>A</sup> | 9.3 (8.0 - 10.6) |  |  |  |
| Mean | 8.4 (7.2 - 9.6) | 12.1 (10.8 - 13.4) |  |  |  |  |
| Inguinal |  |  |  |  |  |  |
| CT | 23.5 (13.8 - 33.3) | 28.8 (20.1 - 37.5) | 26.2 (19.6 - 32.7) <sup>b</sup> | 0.001 | 0.495 | 0.090 |
| HF | 51.6 (41.8 - 61.3) | 39.6 (29.9 - 61.3) | 45.6 (38.7 - 52.5) <sup>a</sup> |  |  |  |
| Mean | 37.5 (30.6 - 44.5) | 34.2 (27.7 - 40.8) |  |  |  |  |
| Epididymal |  |  |  |  |  |  |
| CT | 15.8 (11.1 - 20.5) | 21.1 (17.3 - 25.0) | 18.5 (15.4 - 21.5) | 0.648 | <0.001 | 0.014 |
| HF | 11.0 (6.8 - 15.2) <sup>B</sup> | 27.9 (23.7 - 32.1) <sup>A</sup> | 19.5 (16.5 - 22.4) |  |  |  |
| Mean | 13.4 (10.3 - 16.6) | 25.5 (21.7 - 27.3) |  |  |  |  |
| MDA (nmol/g) |  |  |  |  |  |  |
| Inguinal |  |  |  |  |  |  |
| CT | 3.2 (2.5 - 3.9) | 4.4 (3.7 - 5.2) | 3.8 (3.3 - 4.3) <sup>a</sup> | 0.005 | 0.060 | 0.151 |
| HF | 2.6 (1.9 - 3.3) | 2.8 (2.1 - 3.5) | 2.7 (2.2 - 3.2) <sup>b</sup> |  |  |  |
| Mean | 2.9 (2.4 - 3.4) | 3.6 (3.1 - 4.1) |  |  |  |  |
| Epididymal |  |  |  |  |  |  |
| CT | 3.6 (2.8 - 4.3) | 1.5 (0.7 - 2.3) | 2.5 (2.0 - 3.1) | 0.629 | 0.002 | 0.103 |
| HF | 3.1 (2.3 - 3.8) | 2.3 (1.6 - 3.1) | 2.7 (2.2 - 3.2) |  |  |  |
| Mean | 3.3 (2.8 - 3.8) <sup>A</sup> | 1.9 (1.4 - 2.5) <sup>B</sup> |  |  |  |  |

Values are expressed as mean ± confidence limits. Means differ (P < 0.05); n= 4-7/group. CT: rats fed with control diet (AIN-93G); HF: rats fed with high-fat diet; (+): rats fed with coffee; (-): rats fed without coffee. Capital letters indicate differences between values in the same row, while lowercase letters indicate differences in the same column.

Analysis of lipid oxidation (MDA) in iWAT showed that the HF diet decreased MDA levels compared to that with the CT diet. In the eWAT, coffee consumption reduced MDA concentration compared to that with no coffee (p = 0.002).

### Antioxidant enzyme activity

Coffee consumption significantly reduced CAT activity in BAT (p = 0.004; Table 5). For GST activity, a diet x coffee interaction was exhibited (p = 0.038), with the HF+ diet promoting an increase (p < 0.0001) in GST activity compared to that with the HF diet (p < 0.0001). No difference was observed in the GPx activity after Tukey’s post-hoc adjustment.

**Table 5.**
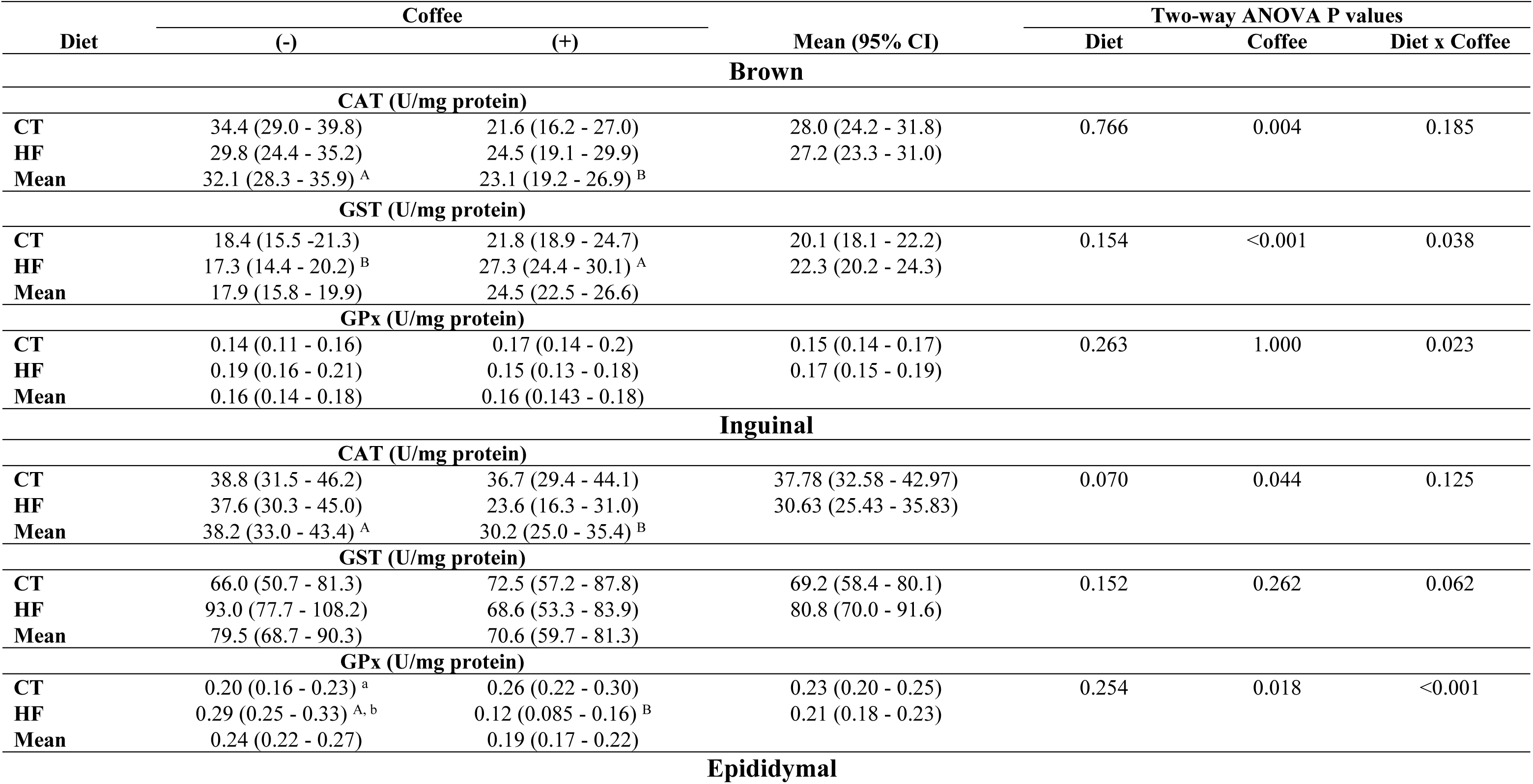
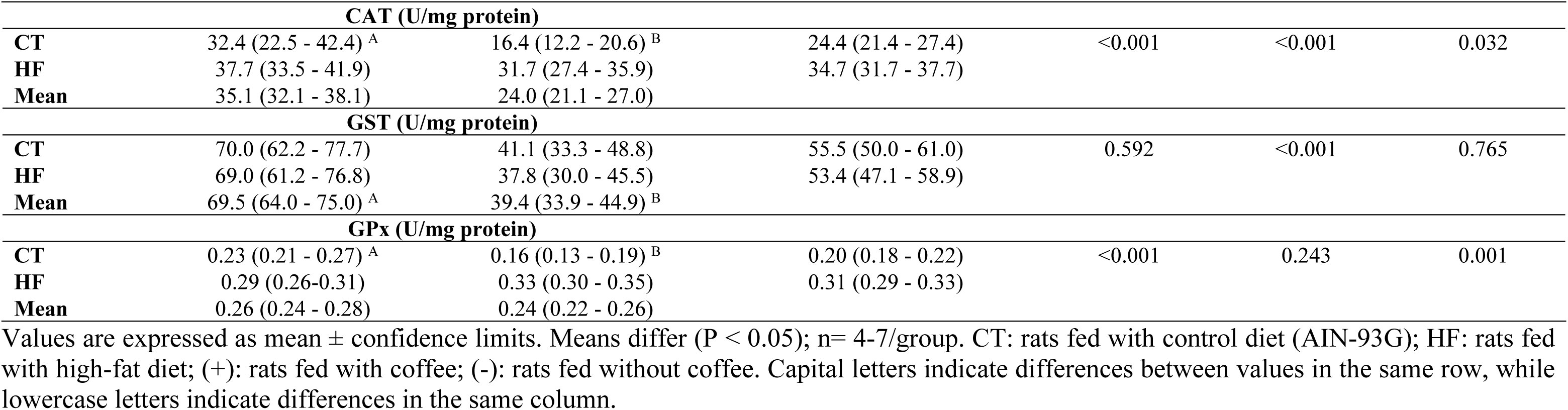
Specific activity of the antioxidant enzymes in adipose tissues of rats treated with control diet (CT) or high-fat diet (HF) with (+) or without (-) coffee.

Similar to that observed in BAT, coffee consumption decreased CAT activity (p = 0.044) compared to that with no coffee in iWAT. A marginal reduction in CAT activity was also observed with a HF diet compared to that with the CT diet (p = 0.070). Regarding GPx activity, a diet x coffee interaction (p < 0.0001) was obtained; the HF+ diet reduced GPx activity compared to that with the HF diet, whereas the HF diet increased GPx activity compared to that with the CT diet (p = 0.004). No treatment effect was observed on GST activity in the iWAT.

Contrary to what was observed in BAT and iWAT, coffee consumption reduced CAT activity in eWAT only when combined with the control diet (CT+), compared to that with the CT diet (p < 0.0001). GST activity was reduced by coffee consumption, regardless of diet type (p < 0.001). Regarding GPx activity, a diet x coffee interaction was identified (p = 0.001), with coffee consumption combined with the control diet (CT+) promoting a decrease in GPx activity compared to that with the CT (p = 0.002).

## Discussion

A HF diet, primarily composed of saturated fatty acids, induced significant alterations in lipid metabolism and adiposity in Wistar rats. Although the total food intake was reduced compared to that of the control group, rats fed a HF diet exhibited greater body weight gain. This paradox is explained by the higher energy density of the diet, which leads to an overall increase in caloric intake despite a lower food volume [35–37]. Levin and Keesey [38] demonstrated that obese rats could adjust their food intake in response to energy demands to regulate weight gain. These findings support our data and are consistent with the current literature on HF diet-induced obesity in rodents.

Marked negative alterations were observed in the histological characteristics of adipose tissues, liver, and lipid profile after consumption of a HF diet for 56 days. There was an increase in the adipocyte area in BAT, iWAT, and eWAT, and the number of unilocular adipocytes in BAT. These findings are consistent with adipocyte hypertrophy, which is a hallmark of obesity. Similarly, Trindade et al. [39] reported an increased fat accumulation in BAT and larger subcutaneous adipocytes were observed in C57BL/6 mice fed a HF diet. Adipocyte hypertrophy is associated with metabolic dysfunction, enhanced lipolysis, and altered adipokine secretion [40,41]. We also observed an increase in hepatic cholesterol and triglyceride levels, and a reduction in the serum LDL/HDL ratio and HDL concentration in rats treated with a HF diet. These data are in accordance with the results obtained in the hepatic histological analysis, as a HF diet increased the microvesicular hypertrophy and liver steatosis scores. These histological changes are characteristic of hepatic steatosis driven by an elevated influx of circulating free fatty acids [42] and consequent lipid accumulation associated with the expansion of intracellular lipid droplets, which store triglycerides and cholesterol esters.

The upregulation of *Vegfa* and *Vegfr2* mRNA levels in BAT suggests that vascularization and consequent blood flow in BAT were induced by the HF diet, which is in accordance with the increased *Ppargc1a and Ucp1* mRNA levels, indicating an improvement in thermogenic capacity. However, this response was not sufficient to completely inhibit BAT whitening in the HF diet group as there was an increase in the area and number of unilocular adipocytes. In iWAT, the increase in *Vegfr2* suggests that a HF diet stimulates angiogenesis in response to the metabolic demands of expanding adipose tissue as a consequence of the high caloric density of the diet. This response was evidenced by higher adipocyte hypertrophy of the iWAT depot compared to that of the eWAT (1.4-fold). Angiogenesis is essential for adipose tissue growth and remodeling, ensuring adequate oxygen and nutrient supply, while facilitating the removal of metabolic byproducts [43].

In contrast, the two key genes involved in *de novo* lipogenesis, *Acaca* and *Fasn,* were downregulated in BATs and eWATs by the HF diet, suggesting a metabolic adaptation to reduce fatty acid synthesis in the context of adipocyte lipid overload. This negative regulation is a well-documented compensatory mechanism in diet-induced obesity models and constitutes a metabolic adaptation to prevent excessive lipid accumulation in adipocytes [44–46]. In addition, in eWAT, the downregulation of *Prkaa1* and *Prkaa2* mRNA levels by the HF diet suggests a reduction in fatty acid uptake by these adipocytes, as AMPK increases glucose and fatty acid uptake in WAT [47]. These results may explain the reduced adipocyte area observed in BAT (1,002.64 µm) and eWAT (3,138.79 µm) compared with that of iWAT (4,482.87 µm). Moreover, in the iWAT, both *Acaca* and *Fasn* mRNA levels were not affected by the HF diet. The upregulation of *Slc2a4* (GLUT4) mRNA levels in the iWAT and eWAT following a HF diet was in accordance with the higher HOMA-IR and HOMA β indices published in our previous study [48]. The high-energy density of the HF diet stimulated glucose uptake, and storage as fatty acids in specialized iWAT and eWAT deposits. Therefore, the HF diet induced insulin resistance, as the increase in *Slc2a4* (GLUT4) mRNA was accompanied by higher insulin secretion [44].

Lipid overload also affects thermogenesis in adipose tissue. The inhibition of the browning capacity of iWAT by HF diet consumption was evidenced by the decrease in *Ucp1* mRNA levels and UCP1 protein content observed in immunohistochemical analysis, which may have contributed to fat accumulation and weight gain. However, *Ucp1* mRNA levels were not affected by the HF diet in the eWAT. Although WAT is the major storage of excess nutrient intake in the form of triglycerides, different WAT depots may not respond similarly to dietary intervention [49]. iWAT exhibits higher susceptibility to browning and greater *Ucp1* expression potential than that of eWAT, which may explain the greater susceptibility of iWAT to whitening under chronic metabolic stress induced by a HF diet [50]. In contrast, in BAT, *Ucp1* and *Ppargc1a* mRNA levels increased, suggesting that the HF diet enhanced its thermogenic capacity. The greater availability of fatty acids in the HF diet may have enhanced *Ucp1* mRNA levels [51] in an attempt to dissipate excess energy [52].

In HF diet-induced obesity models, inflammatory processes may modulate thermogenic activity. To explore this possibility, we quantified the mRNA levels of *Il10* and *Mcp1* in adipose tissues. However, the role of the anti-inflammatory cytokine, IL-10, in BAT activity remains controversial. IL-10 is essential for BAT function and thermogenesis because its deficiency leads to marked mitochondrial dysfunction and impaired cold tolerance [53]. Conversely, other evidence suggests that IL-10 may either have no significant impact or even exert a suppressive effect on thermogenesis and energy expenditure [54]. In our study, a HF diet induced the adrenergic activation of BAT, as demonstrated by the upregulation of thermogenesis-related genes (*Ppargc1a*, *Ucp1*, *Vegfa*, and *Vegfr2*) and increased *Il10* mRNA expression. These findings suggest that a possible compensatory anti-inflammatory response contributes to BAT activation in diet-induced obesity.

Obesity is strongly associated with oxidative damage to biomolecules [55]. Long et al. [56] suggested that the high content of polyunsaturated fatty acids (PUFA) in adipose tissue results in lipid peroxidation and lipid peroxides may induce protein carbonylation. In the present study, a HF diet reduced lipid peroxidation (MDA) but increased the carbonyl content in iWAT. These results may be related to the low PUFA concentration in the lard and high GPx activity compared to those with the CT diet, as GPx catalyzes the decomposition of H_2_O_2_ and organic hydroperoxides, using reduced glutathione as a cofactor. The two-fold increase in protein carbonylation in iWAT was consistent with previous reports on HF-diet models [57]. Galinier et al. [58] reported lower MDA concentrations in the iWAT of obese rats than that in lean rats. The authors demonstrated that fat accumulation was associated with higher concentrations of hydrophilic antioxidants, which can regenerate lipophilic antioxidants, such as tocopherol radicals. Therefore, the authors suggested a role for hydrophilic antioxidants in lipid peroxidation inhibition. However, in our study, despite the high metabolic rate of eWAT in the HF diet-treated rats, as evidenced by the reduced adipocyte area and downregulation of *Acaca* and *Fasn* mRNA levels compared with those in iWAT, no significant differences in the redox response were observed in eWAT.

Coffee contains a variety of phytochemicals with recognized antioxidant and anti-inflammatory properties [59,60], which may attenuate the whitening of WATs and activate BAT activity by modulating lipid metabolism and adiposity. In the present study, coffee consumption combined with a HF diet did not affect overall body weight gain compared to that with the HF diet alone. However, the combination of coffee and HF diet appeared to enhance BAT activation, as evidenced by a reduction in the adipocyte area and low number of unilocular adipocytes. This result was in accordance with the downregulation of *Acaca* and *Fasn* mRNA levels observed after coffee consumption in BAT. The first gene encodes the enzyme acetyl-CoA carboxylase, which produces malonyl-CoA and inhibits the entry of fatty acids into the mitochondria and, consequently, β-oxidation. The second gene encodes fatty acid synthase, the enzyme that catalyzes the synthesis of fatty acids. The redox response was linked to BAT remodeling during thermogenesis induction, in which chronic treatment (40–45 days) decreases reactive oxygen species (ROS) production and enzymatic antioxidant defenses [61]. Therefore, the reduction in catalase activity in BAT by coffee consumption, regardless of diet type, reinforces the hypothesis of inhibition of BAT whitening by coffee, as uncoupling has been associated with lower ROS production and antioxidant enzymatic activity. However, no effect of coffee was observed in the thermogenic gene, *Ppargc1a*, *Ucp1*, *Prdm16*, *Vegfa, and Vegfr2,* mRNA levels in BAT, regardless of diet type.

In the iWAT, coffee reduced the whitening effects typically induced by a HF diet, as the adipocyte area was reduced when coffee was administered alongside the HF diet. Additionally, an increase in the number of multilocular adipocytes was observed regardless of diet type. Therefore, coffee was able to inhibit the whitening process in the iWAT, even in rats fed the CT diet. These results are in accordance with the upregulation of thermogenic and angiogenic genes, *Prdm16* and *Vegfr2,* observed in response to coffee consumption, independent of the diet type. This hypothesis is supported by the immunohistochemical results, which demonstrated that UCP1 protein levels in iWAT were increased by coffee consumption, regardless of the diet type. The increase in protein levels, but not mRNA levels UCP1 in iWAT, suggests that coffee compounds modulate UCP1 at the translational or post-translational level. Considering that uncoupling leads to a reduction in ROS production [61,62], the decrease in CAT activity by coffee, regardless of diet type, and GPx activity when coffee was combined with the HF diet, supports the hypothesis that coffee inhibits the whitening of iWAT. The upregulation of *Fasn* mRNA levels by coffee occurred only when combined with the CT diet in the iWAT and may be related to the high availability of glucose in this diet compared to that with the HF diet. Previously, the HF diet downregulated *Slc2a2* and *Slc5a5* mRNA levels in the intestine, and coffee decreased intestinal α-glucosidase activity compared to the HF diet alone [48]. The reduction in *Mcp1* mRNA levels in iWAT following coffee consumption, in conjunction with the CT diet, suggests an improvement in the anti-inflammatory response, given that this chemokine is known to mediate macrophage recruitment into adipose tissue. The resulting anti-inflammatory phenotype promotes a metabolically favorable environment that supports beiging of WAT [4].

Coffee consumption also effectively suppressed the whitening characteristics in eWAT, independent of diet type. This effect was evidenced by an increased number of multilocular adipocytes and *Prdm16* mRNA levels, which are key regulators of adipose tissue browning. Moreover, *Fasn* mRNA levels, which are responsible for *de novo* fatty acid synthesis, were reduced compared to those in their non-coffee counterparts. These results suggest that coffee consumption promotes enhanced thermogenesis and the consequent attenuation of adiposity. Upregulation of *Prdm16* mRNA supports beige adipocyte differentiation, even in the absence of *Ucp1* mRNA modulation, indicating UCP1-independent thermogenesis or transitional paucilocular adipocytes not yet expressing *Ucp1* [49,63]. Similarly, Gamboa-Gómez et al. [64] observed a decrease in adipocyte volume following the administration of unroasted and roasted coffee in obese rats. The coffee thermogenic effect in the eWAT occurred even in the control condition, as *Ppargc1a* mRNA levels increased and *Acaca* mRNA levels, which inhibit lipolysis, decreased in CT+ rats compared with those in CT rats. In the control diet group, coffee also inhibited CAT, GPx activity, and MDA, suggesting that uncoupling was activated in eWAT, as this process was associated with a decrease in ROS production and consequent decrease in antioxidant enzymatic activity.

The increasing prevalence of obesity is paralleled by an increase in the incidence of NAFLD. Certain diterpenes present in coffee, such as cafestol and kahweol, increase cholesterol levels by reducing the excretion of bile acids and neutral sterols [65,66]. Although the coffee used in this study was prepared with boiled water and paper filters, which significantly reduced diterpene content, histological analysis of the liver tissue revealed that the CT+ diet led to high scores for NAFLD-related parameters, including microvesicular steatosis, hepatocellular hypertrophy, and overall liver steatosis. In line with these findings, coffee consumption promoted the accumulation of triglycerides and cholesterol in the hepatic tissues, regardless of the diet type. These results suggest that under a controlled diet, coffee may contribute to hepatic lipid accumulation in the form of numerous small lipid vesicles within hepatocytes. However, as observed in several studies [67,68], coffee consumption combined with a high-fat diet ameliorated hypertrophy and liver steatosis scores. This response may be related to the high cholesterol content of lard used to formulate the diet, since an initial increase in intracellular cholesterol levels may lead to a reduction in LDL receptors to control cell cholesterol concentration. The increase in serum LDL/HDL ratio with coffee consumption combined with a high-fat diet reinforces this hypothesis. Senftinger et al. [69] observed in a cross-sectional analysis of the population-based Hamburg City Health Study that high coffee consumption was correlated with high circulating LDL content, suggesting that the diterpenes present in coffee reduce LDL receptor activity and intracellular cholesterol concentration. Additionally, some phenolic compounds found in coffee, such as chlorogenic acids, have been associated with the activation of metabolic pathways that favor hepatic fatty acid β-oxidation and suppress lipogenesis [70–73].

## Conclusions

The consumption of coffee by rats at doses comparable to the usual intake of the Brazilian population partially attenuated the whitening process in BAT, likely through the modulation of lipogenic and lipolytic gene expression. In iWAT, coffee exhibited a more pronounced effect than that of eWAT by enhancing the expression of thermogenic and angiogenic genes, thereby reducing the whitening features, regardless of whether animals were maintained on a control or high-fat diet. Collectively, these findings provide insight into the molecular mechanisms by which coffee may contribute to obesity management.

## Acknowledgments

We would like to thank Editage (www.editage.com.br) for English language editing.

## Notes

### Competing Interest Statement

The authors have declared no competing interest.

## References

1. Zhang H, Zhou XD, Shapiro MD, Lip GYH, Tilg H, Valenti L, et al. Global burden of metabolic diseases, 1990–2021. Metabolism. 2024;160: 155999. doi:10.1016/j.metabol.2024.155999.

2. Swinburn BA, Kraak VI, Allender S, Atkins VJ, Baker PI, Bogard JR, et al. The Global Syndemic of Obesity, Undernutrition, and Climate Change: The Lancet Commission report. The Lancet. 2019;393: 791–846. doi:10.1016/S0140-6736(18)32822-8.

3. Machado SA, Pasquarelli-do-Nascimento G, da Silva DSS, Farias GR, de Oliveira Santos I, Baptista LB, et al. Browning of the white adipose tissue regulation: new insights into nutritional and metabolic relevance in health and diseases. Nutrition & Metabolism 2022 19:1. 2022;19: 1–27. doi:10.1186/S12986-022-00694-0.

4. Villarroya F, Cereijo R, Gavaldà-Navarro A, Villarroya J, Giralt M. Inflammation of brown/beige adipose tissues in obesity and metabolic disease. J Intern Med. 2018;284: 492–504. doi:10.1111/joim.12803.

5. Kuryłowicz A, Puzianowska-Kuźnicka M. Induction of Adipose Tissue Browning as a Strategy to Combat Obesity. Int J Mol Sci. 2020;21: 6241. doi:10.3390/ijms21176241.

6. Chiu YH, Chou WL, Ko MC, Liao JC, Huang TH. Curcumin mitigates obesity-driven dysbiosis and liver steatosis while promoting browning and thermogenesis in white adipose tissue of high-fat diet-fed mice. J Nutr Biochem. 2025;143: 109920. doi:10.1016/j.jnutbio.2025.109920.

7. Li X, Yao Z, Qi X, Cui JL, Zhou Y, Tan Y, et al. Naringin ameliorates obesity via stimulating adipose thermogenesis and browning, and modulating gut microbiota in diet-induced obese mice. Curr Res Food Sci. 2024;8: 100683. doi:10.1016/j.crfs.2024.100683.

8. Evans BA, Merlin J, Bengtsson T, Hutchinson DS. Adrenoceptors in white, brown, and brite adipocytes. Br J Pharmacol. 2019;176: 2416–2432. doi:10.1111/bph.14631.

9. Lidell ME, Betz MJ, Enerbäck S. Brown adipose tissue and its therapeutic potential. J Intern Med. 2014;276: 364–377. doi:10.1111/joim.12255.

10. Chouchani ET, Kazak L, Spiegelman BM. New Advances in Adaptive Thermogenesis: UCP1 and Beyond. Cell Metab. 2019;29: 27–37. doi:10.1016/j.cmet.2018.11.002.

11. Fu M, Yoon KS, Ha J, Kang I, Choe W. Crosstalk Between Antioxidants and Adipogenesis: Mechanistic Pathways and Their Roles in Metabolic Health. Antioxidants. 2025;14: 203. doi:10.3390/antiox14020203.

12. Zheng Y, Yang N, Pang Y, Gong Y, Yang H, Ding W, et al. Mitochondria-associated regulation in adipose tissues and potential reagents for obesity intervention. Front Endocrinol (Lausanne). 2023;14: 1132342. doi:10.3389/fendo.2023.1132342.

13. Chouchani ET, Kajimura S. Metabolic adaptation and maladaptation in adipose tissue. Nature Metabolism 2019 1:2. 2019;1: 189–200. doi:10.1038/s42255-018-0021-8.

14. Liu X, Zhang Z, Song Y, Xie H, Dong M. An update on brown adipose tissue and obesity intervention: Function, regulation and therapeutic implications. Front Endocrinol (Lausanne). 2023;13: 1065263. doi:10.3389/fendo.2022.1065263.

15. Sakers A, De Siqueira MK, Seale P, Villanueva CJ. Adipose-tissue plasticity in health and disease. Cell. 2022;185: 419–446. doi:10.1016/j.cell.2021.12.016.

16. Kawai T, Autieri M V., Scalia R. Adipose tissue inflammation and metabolic dysfunction in obesity. Am J Physiol Cell Physiol. 2021;320: C375–C391. doi:10.1152/ajpcell.00379.2020.

17. Stefanello N, Spanevello RM, Passamonti S, Porciúncula L, Bonan CD, Olabiyi AA, et al. Coffee, caffeine, chlorogenic acid, and the purinergic system. Food and Chemical Toxicology. 2019;123: 298–313. doi:10.1016/j.fct.2018.10.005.

18. Higdon J V., Frei B. Coffee and health: A review of recent human research. Crit Rev Food Sci Nutr. 2006;46: 101–123. doi:10.1080/10408390500400009.

19. Kobylińska Z, Biesiadecki M, Kuna E, Galiniak S, Mołoń M. Coffee as a Source of Antioxidants and an Elixir of Youth. Antioxidants 2025, Vol 14, Page 285. 2025;14: 285. doi:10.3390/antiox14030285.

20. Farias-Pereira R, Park CS, Park Y. Mechanisms of action of coffee bioactive components on lipid metabolism. Food Sci Biotechnol. 2019;28: 1287. doi:10.1007/S10068-019-00662-0.

21. Salazar J, Cano C, Pérez JL, Castro A, Díaz MP, Garrido B, et al. Role of Dietary Polyphenols in Adipose Tissue Browning: A Narrative Review. Curr Pharm Des. 2020;26: 4444–4460. doi:10.2174/1381612826666200701211422.

22. ABIC. Métodos de preparo. Rio de Janeiro: Associação Brasileira da Indústria de Café; 2021. Available: https://www.abic.com.br/tudo-de-cafe/metodos-de-preparo/.

23. Sousa AG, Da Costa THM. Usual coffee intake in Brazil: Results from the National Dietary Survey 2008-9. British Journal of Nutrition. 2015;113: 1615–1620. doi:10.1017/S0007114515000835.

24. National Research Council (US) Committee for the Update of the Guide for the Care and Use of Laboratory Animals. Guide for the care and use of laboratory animals. 8th ed. Washington (DC): National Academies Press; 2011. doi: 10.17226/12910.

25. Reeves PG, Nielsen FH, Fahey GC. AIN-93 purified diets for laboratory rodents: Final report of the American Institute of Nutrition ad hoc writing committee on the reformulation of the AIN-76A rodent diet. Journal of Nutrition. 1993;123: 1939–1951. doi:10.1093/jn/123.11.1939.

26. Vieira VJ, Valentine RJ, Wilund KR, Woods JA. Effects of diet and exercise on metabolic disturbances in high-fat diet-fed mice. Cytokine. 2009;46: 339–345. doi:10.1016/j.cyto.2009.03.006.

27. Liang W, Menke AL, Driessen A, Koek GH, Lindeman JH, Stoop R, et al. Establishment of a General NAFLD Scoring System for Rodent Models and Comparison to Human Liver Pathology. PLoS One. 2014;9: e115922. doi:10.1371/journal.pone.0115922.

28. Mesquita CS, Oliveira R, Bento F, Geraldo D, Rodrigues J V., Marcos JC. Simplified 2,4-dinitrophenylhydrazine spectrophotometric assay for quantification of carbonyls in oxidized proteins. Anal Biochem. 2014;458: 69–71. doi:10.1016/j.ab.2014.04.034.

29. De-Souza MT, Brígido MDM, Maranhão AQ. Técnicas básicas em biologia molecular. 2a. UnB, Editora. Brasília; 2016.

30. Livak KJ, Schmittgen TD. Analysis of Relative Gene Expression Data Using Real-Time Quantitative PCR and the 2−ΔΔCT Method. Methods. 2001;25: 402–408. doi:10.1006/meth.2001.1262.

31. Hartree EF. Determination of protein: A modification of the lowry method that gives a linear photometric response. Anal Biochem. 1972;48: 422–427. doi:10.1016/0003-2697(72)90094-2.

32. Candan N, Tuzmen N. Very rapid quantification of malondialdehyde (MDA) in rat brain exposed to lead, aluminium and phenolic antioxidants by high-performance liquid chromatography-fluorescence detection. Neurotoxicology. 2008;29: 708–713. doi:10.1016/j.neuro.2008.04.012.

33. Joanisse DR, Storey KB. Oxidative damage and antioxidants in Rana sylvatica, the freeze-tolerant wood frog. Am J Physiol.1996 Sep;271(3 Pt 2):R545-53. doi:10.1152/ajpregu.1996.271.3.r545.

34. Habig WH, Jakoby WB. Glutathione S-Transferases (Rat and Human). Methods Enzymol. 1981;77: 218–231. doi:10.1016/S0076-6879(81)77029-0.

35. Di Majo D, Sardo P, Giglia G, Di Liberto V, Zummo FP, Zizzo MG, et al. Correlation of Metabolic Syndrome with Redox Homeostasis Biomarkers: Evidence from High-Fat Diet Model in Wistar Rats. Antioxidants. 2022;12: 89. doi:10.3390/antiox12010089.

36. Hu S, Wang L, Yang D, Li L, Togo J, Wu Y, et al. Dietary Fat, but Not Protein or Carbohydrate, Regulates Energy Intake and Causes Adiposity in Mice. Cell Metab. 2018;28: 415–431.e4. doi:10.1016/j.cmet.2018.06.010.

37. White PAS, Cercato LM, Araújo JMD, Souza LA, Soares AF, Barbosa APO, et al. Modelo de obesidade induzida por dieta hiperlipídica e associada à resistência à ação da insulina e intolerância à glicose. Arquivos Brasileiros de Endocrinologia & Metabologia. 2013;57: 339–345. doi:10.1590/S0004-27302013000500002.

38. Levin BE, Keesey RE. Defense of differing body weight set points in diet-induced obese and resistant rats. Am J Physiol Regul Integr Comp Physiol. 1998;274. doi:10.1152/ajpregu.1998.274.2.R412.

39. Trindade PL, Martins FF, dos Ramos Soares E, Bernardes EM, Vardiero F, de Castro Resende A, et al. Polyphenol-rich jaboticaba (Myrciaria jaboticaba) peel and seed powder induces browning of subcutaneous white adipose tissue and improves metabolic status in high-fat-fed mice. J Funct Foods. 2022;97: 105238. doi:10.1016/j.jff.2022.105238.

40. Mallick R, Basak S, Das RK, Banerjee A, Paul S, Pathak S, et al. Fatty Acids and their Proteins in Adipose Tissue Inflammation. Cell Biochemistry and Biophysics 2023 82:1. 2023;82: 35–51. doi:10.1007/S12013-023-01185-6.

41. Li Q, Spalding KL. The regulation of adipocyte growth in white adipose tissue. Front Cell Dev Biol. 2022;10: 1003219. doi:10.3389/fcell.2022.1003219.

42. Heeren J, Scheja L. Metabolic-associated fatty liver disease and lipoprotein metabolism. Mol Metab. 2021;50: 101238. doi:10.1016/j.molmet.2021.101238.

43. Sreekumar S, Gangaraj KP, Kiran MS. Modulation of angiogenic switch in reprogramming browning and lipid metabolism in white adipocytes. Biochimica et Biophysica Acta (BBA) - Molecular and Cell Biology of Lipids. 2024;1869: 159423. doi:10.1016/j.bbalip.2023.159423.

44. Smith U, Kahn BB. Adipose tissue regulates insulin sensitivity: role of adipogenesis, de novo lipogenesis and novel lipids. J Intern Med. 2016;280: 465. doi:10.1111/joim.12540.

45. Marcelino H, Veyrat-Durebex C, Summermatter S, Sarafian D, Miles-Chan J, Arsenijevic D, et al. A Role for Adipose Tissue De Novo Lipogenesis in Glucose Homeostasis During Catch-up Growth: A Randle Cycle Favoring Fat Storage. Diabetes. 2013;62: 362–372. doi:10.2337/DB12-0255.

46. Mottillo EP, Balasubramanian P, Lee YH, Weng C, Kershaw EE, Granneman JG. Coupling of lipolysis and de novo lipogenesis in brown, beige, and white adipose tissues during chronic β3-adrenergic receptor activation. J Lipid Res. 2014;55: 2276. doi:10.1194/jlr.M050005.

47. Göransson O, Kopietz F, Rider MH. Metabolic control by AMPK in white adipose tissue. Trends in Endocrinology & Metabolism. 2023;34: 704–717. doi:10.1016/j.tem.2023.08.011.

48. Ramos LV, da Costa THM, Arruda SF. The effect of coffee consumption on glucose homeostasis and redox-inflammatory responses in high-fat diet-induced obese rats. J Nutr Biochem. 2022;100: 108881. doi:10.1016/j.jnutbio.2021.108881.

49. Duerre DJ, Galmozzi A. Deconstructing Adipose Tissue Heterogeneity One Cell at a Time. Front Endocrinol (Lausanne). 2022;13: 847291. doi:10.3389/fendo.2022.847291.

50. Chi J, Wu Z, Choi CHJ, Nguyen L, Tegegne S, Ackerman SE, et al. Three-Dimensional Adipose Tissue Imaging Reveals Regional Variation in Beige Fat Biogenesis and PRDM16-Dependent Sympathetic Neurite Density. Cell Metab. 2018;27: 226–236.e3. doi:10.1016/j.cmet.2017.12.011.

51. Fromme T, Klingenspor M. Uncoupling protein 1 expression and high-fat diets. Am J Physiol Regul Integr Comp Physiol. 2011;300: 1–8. doi:10.1152/ajpregu.00411.2010.

52. Ahluwalia R, Luijten IHN, Sousa-Filho CPB, Braz GRF, Petrovic N, Shabalina IG, et al. The choice of diet is determinative for the manifestation of UCP1-dependent diet-induced thermogenesis. Am J Physiol Endocrinol Metab. 2025;328: E653–E660. doi:10.1152/ajpendo.00038.2025.

53. de-Lima-Júnior JC, Souza GF, Moura-Assis A, Gaspar RS, Gaspar JM, Rocha AL, et al. Abnormal brown adipose tissue mitochondrial structure and function in IL10 deficiency. Ebio Medicine. 2019; 39: 436–447. doi:10.1016/j.ebiom.2018.11.041.

54. Rajbhandari P, Thomas BJ, Feng AC, Hong C, Wang J, Vergnes L, et al. IL-10 Signaling Remodels Adipose Chromatin Architecture to Limit Thermogenesis and Energy Expenditure. Cell. 2018;172: 218–233.e17. doi:10.1016/j.cell.2017.11.019.

55. Matsuda M, Shimomura I. Increased oxidative stress in obesity: Implications for metabolic syndrome, diabetes, hypertension, dyslipidemia, atherosclerosis, and cancer. Obes Res Clin Pract. 2013;7: e330–e341. doi:10.1016/j.orcp.2013.05.004.

56. Long EK, Olson DM, Bernlohr DA. High-fat diet induces changes in adipose tissue trans-4-oxo-2-nonenal and trans-4-hydroxy-2-nonenal levels in a depot-specific manner. Free Radic Biol Med. 2013;63: 390–398. doi:10.1016/j.freeradbiomed.2013.05.030.

57. Grimsrud PA, Picklo MJ, Griffin TJ, Bernlohr DA. Carbonylation of Adipose Proteins in Obesity and Insulin Resistance: Identification of Adipocyte Fatty Acid-binding Protein as a Cellular Target of 4-Hydroxynonenal. Molecular & Cellular Proteomics. 2007;6: 624–637. doi:10.1074/MCP.M600120-MCP200.

58. Galinier A, Carrière A, Fernandez Y, Carpéné C, André M, Caspar-Bauguil S, et al. Adipose Tissue Proadipogenic Redox Changes in Obesity. Journal of Biological Chemistry. 2006;281: 12682–12687. doi:10.1074/jbc.M506949200.

59. León D, Medina S, Londoño-Londoño J, Jiménez-Cartagena C, Ferreres F, Gil-Izquierdo A. Anti-inflammatory Activity of Coffee. In: Farah A, editor. Coffee: Consumption and Health Implications. Royal Society of Chemistry; 2019. pp. 57–74. doi:10.1039/9781788015028-00057.

60. del Castillo MD, Iriondo-DeHond A, Fernandez-Gomez B, Martinez-Saez N, Rebollo-Hernanz M, Martín-Cabrejas MA, et al. Coffee Antioxidants in Chronic Diseases. In: Farah A, editor. Coffee. Royal Society of Chemistry; 2019. pp. 20–56. doi:10.1039/9781788015028-00020.

61. Petrović V, Buzadžić B, Korać A, Vasilijević A, Janković A, Korać B. Free radical equilibrium in interscapular brown adipose tissue: Relationship between metabolic profile and antioxidative defense. Comparative Biochemistry and Physiology Part C: Toxicology & Pharmacology. 2006;142: 60–65. doi:10.1016/j.cbpc.2005.10.004.

62. Skulachev VP. Snizhenie vnizhenie vnutrikletochnoĭ kontsentratsii O2 kak osobaia dykhatel’nykh sistem kletki [Decrease in the intracellular concentration of O2 as a special function of the cellular respiratory system]. Biokhimiia. 1994;59: 1910–2.

63. Jiang N, Yang M, Han Y, Zhao H, Sun L. PRDM16 Regulating Adipocyte Transformation and Thermogenesis: A Promising Therapeutic Target for Obesity and Diabetes. Front Pharmacol. 2022;13: 870250. doi:10.3389/fphar.2022.870250.

64. Gamboa-Gómez CI, Morales-Castro J, Barragan-Zuñiga J, Herrera MD, Zamilpa-Álvarez A, Gónzalez JL, et al. Influence of coffee roasting degree on antioxidant and metabolic parameters: Comprehensive in vitro and in vivo analysis. Curr Res Food Sci. 2024;9: 100861. doi:10.1016/j.crfs.2024.100861.

65. O’Keefe JH, Bhatti SK, Patil HR, Dinicolantonio JJ, Lucan SC, Lavie CJ. Effects of Habitual Coffee Consumption on Cardiometabolic Disease, Cardiovascular Health, and All-Cause Mortality. J Am Coll Cardiol. 2013;62: 1043–1051. doi:10.1016/j.jacc.2013.06.035.

66. Jee SH, He J, Appel LJ, Whelton PK, Suh I, Klag MJ. Coffee consumption and serum lipids: A meta-analysis of randomized controlled clinical trials. Am J Epidemiol. 2001;153: 353–362. doi:10.1093/aje/153.4.353.

67. Abrahão SA, Pereira RGFA, de Sousa RV, Lima AR, Crema GP, Barros BS. Influence of Coffee Brew in Metabolic Syndrome and Type 2 Diabetes. Plant Foods for Human Nutrition. 2013;68: 184–189. doi:10.1007/s11130-013-0355-z.

68. Maki C, Funakoshi-Tago M, Aoyagi R, Ueda F, Kimura M, Kobata K, et al. Coffee extract inhibits adipogenesis in 3T3-L1 preadipocyes by interrupting insulin signaling through the downregulation of IRS1. PLoS One. 2017;12: e0173264. doi:10.1371/journal.pone.0173264.

69. Senftinger J, Nikorowitsch J, Borof K, Ojeda F, Aarabi G, Beikler T, et al. Coffee consumption and associations with blood pressure, LDL-cholesterol and echocardiographic measures in the general population. Scientific Reports 2023 13:1. 2023;13: 1–9. doi:10.1038/s41598-023-31857-5.

70. Zhang SJ, Li YF, Wang GE, Tan RR, Tsoi B, Mao GW, et al. Caffeine ameliorates high energy diet-induced hepatic steatosis: sirtuin 3 acts as a bridge in the lipid metabolism pathway. Food Funct. 2015;6: 2578–2587. doi:10.1039/C5FO00247H.

71. Murase T, Misawa K, Minegishi Y, Aoki M, Ominami H, Suzuki Y, et al. Coffee polyphenols suppress diet-induced body fat accumulation by downregulating SREBP-1c and related molecules in C57BL/6J mice. Am J Physiol Endocrinol Metab. 2011;300: 122–133. doi:10.1152/ajpendo.00441.2010.

72. Miao H, Ouyang H, Guo Q, Wei M, Lu B, Kai G, et al. Chlorogenic acid alleviated liver fibrosis in methionine and choline deficient diet-induced nonalcoholic steatohepatitis in mice and its mechanism. J Nutr Biochem. 2022;106: 109020. doi:10.1016/j.jnutbio.2022.109020.

73. Shi A, Li T, Zheng Y, Song Y, Wang H, Wang N, et al. Chlorogenic Acid Improves NAFLD by Regulating gut Microbiota and GLP-1. Front Pharmacol. 2021;12: 693048. doi:10.3389/fphar.2021.693048.

